# Organoid co-culture model of the cycling human endometrium in a fully-defined synthetic extracellular matrix reveals epithelial-stromal crosstalk

**DOI:** 10.1101/2021.09.30.462577

**Authors:** Juan S. Gnecco, Alexander Brown, Kira Buttrey, Clara Ives, Brittany A. Goods, Lauren Baugh, Victor Hernandez-Gordillo, Megan Loring, Keith Isaacson, Linda G. Griffith

**Affiliations:** Center for Gynepathology Research, Massachusetts Institute of Technology, 77 Massachusetts Ave, Cambridge MA, USA 02139; Department of Biological Engineering, Massachusetts Institute of Technology, 77 Massachusetts Ave, Cambridge MA, USA 02139; Thayer School of Engineering at Dartmouth College, Hanover, NH.; Newton Wellesley Hospital, Newton, MA USA 02115.

## Abstract

The human endometrium undergoes recurring cycles of growth, differentiation, and breakdown in response to sex hormones. Dysregulation of epithelial-stromal communication during hormone cycles is linked to myriad gynecological disorders for which treatments remain inadequate. Here, we describe a completely defined, synthetic extracellular matrix that enables co-culture of human endometrial epithelial and stromal cells in a manner that captures healthy and disease states across a simulated menstrual cycle. We parsed cycle-dependent endometrial integrin expression and matrix composition to define candidate cell-matrix interaction cues for inclusion in a polyethylene glycol (PEG)-based hydrogel crosslinked with matrix metalloproteinase-labile peptides. We semi-empirically screened a parameter space of biophysical and molecular features representative of the endometrium to define compositions suitable for hormone-driven expansion and differentiation of epithelial organoids, stromal cells, and co-cultures of the two cell types. Each cell type exhibited characteristic morphological and molecular responses to hormone changes when co-encapsulated in hydrogels tuned to a stiffness regime similar to the native tissue and functionalized with a collagen-derived adhesion peptide (GFOGER) and a fibronectin-derived peptide (PHSRN-K-RGD). Analysis of cell-cell crosstalk during IL-1β-induced inflammation revealed dysregulation of epithelial proliferation mediated by stromal cells. Altogether, we demonstrate the development of a fully synthetic matrix to sustain the dynamic changes of the endometrial microenvironment and support its applications to understand menstrual health and endometriotic diseases.

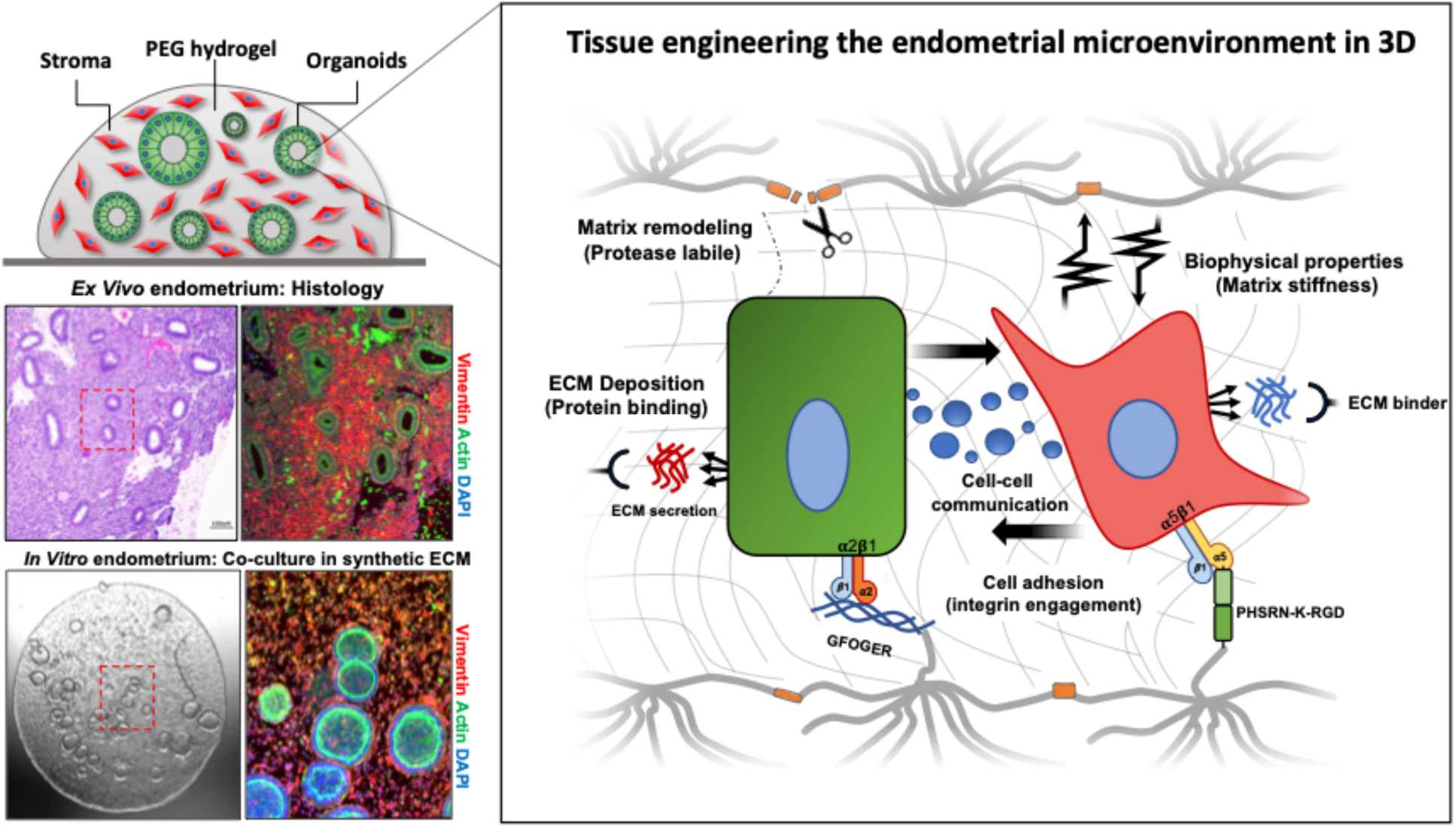

## 1. Introduction

The endometrium is the mucosal lining of the uterus in which the establishment and maintenance of pregnancy occurs^1^. It is a highly dynamic tissue that undergoes spatial and temporal changes in response to endocrine signaling by the ovarian sex hormones estradiol (E2) and progesterone (P4). Cyclical changes in these hormones dictate the timing and functional capabilities of the endometrium to support nidation by driving cell-specific morphological and biochemical processes^2^. Histologically, the endometrium is primarily composed of the hormone-responsive epithelium and specialized reticular fibroblasts (stromal cells) embedded in a dynamic extracellular matrix (ECM).^1, 3, 4^ Endocrine-mediated reproductive function is mediated through both direct and indirect mechanisms governed by the crosstalk between these cell populations.^5–7^ The idealized 28-day human menstrual cycle is characterized by three distinct phases in which the endometrium undergoes E2-mediated tissue regeneration (the ‘proliferative phase’) followed by a 14-day P4-dominant phase (the ‘secretory phase’) that governs the differentiation processes necessary for the successful establishment of pregnancy (Fig 1A). In the absence of embryo implantation, a sharp withdrawal of hormones triggers a cascade of inflammatory processes that result in the shedding of the endometrial tissue (the ‘menstrual phase’) after which the cycle initiates again.^1, 3^ Despite the fundamental roles E2 and P4 play in maintaining reproductive functions, a thorough understanding of the cellular mechanisms driving these processes remains elusive due in part to a lack of physiological models that recapitulate the complex multi-cellular human condition.^1, 8^

**Figure 1.**
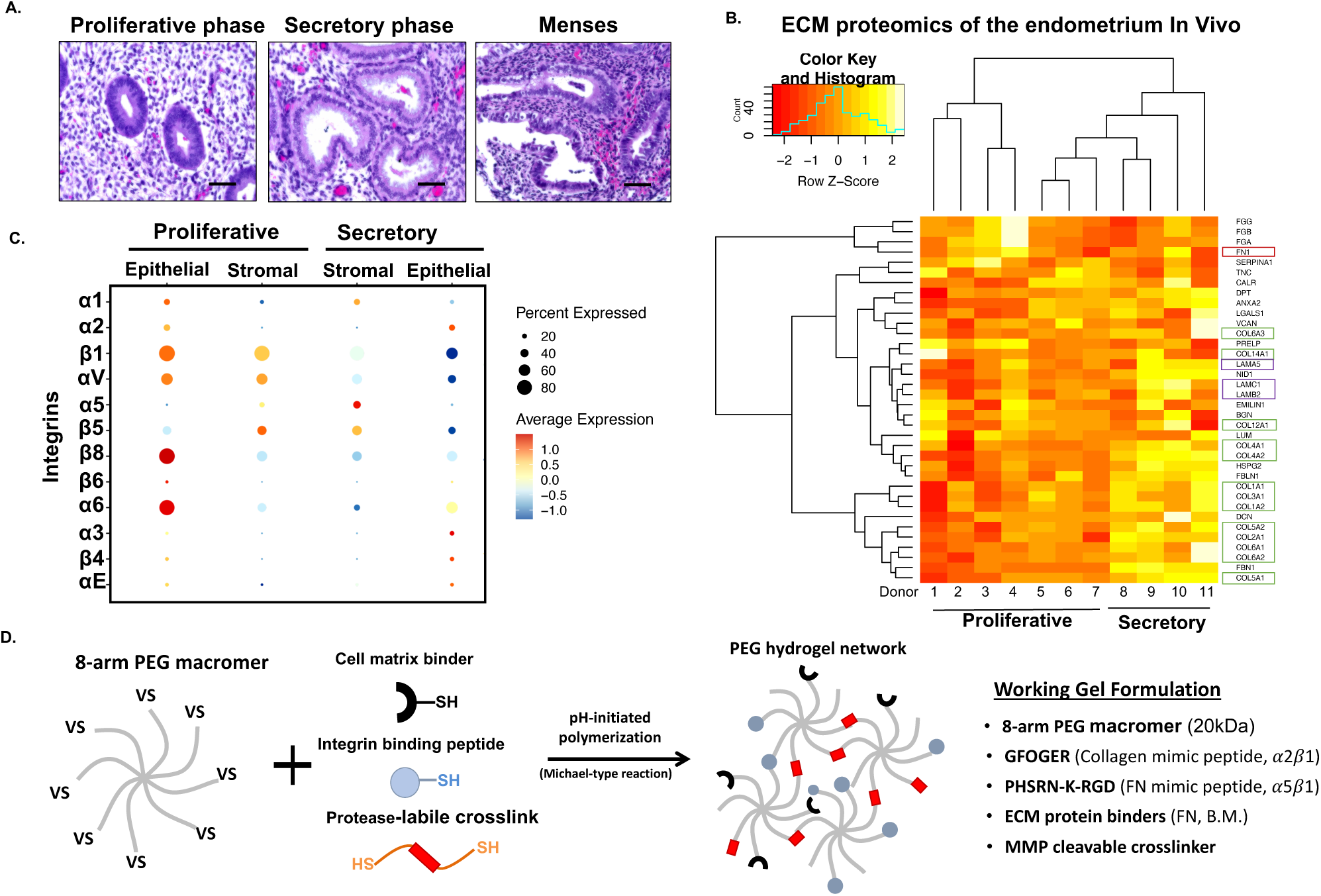
A tissue-inspired multi-omic approach elucidates the design parameters to engineer a fully synthetic PEG hydrogel for the endometrium. (A) Representative histological hematoxylin and eosin (H&E) stained samples of the menstrual cycle phases. (B) Heatmap of proteomic analysis of the human endometrium showing ECM proteins (matrisome) across the menstrual cycle (n=11 donors). Box colors indicated ECM protein associations: green corresponds to collagens; red to fibronectin; and purple to laminins. (C) Dot plot analysis of specific integrins expression from single cell RNA-seq data (n = 6 donors, 6,659 cells). Examination of stromal and epithelial component integrin expression across the menstrual cycle (proliferative and secretory phases). Circle size and color indicate P-value and variation about the means of the average expression value for each integrin chain in all cells (D) Schematic of the overall strategy to design and fabricate a synthetic ECM for the endometrium. 8-arm 20kDa PEG macromers are functionalized with cell matrix binders (FN, B.M.), integrin adhesion peptides, and polymerized with a protease-labile crosslinker. Descriptive outline of the key peptides utilized in the synthetic ECM formulation.

Experimental models capable of recapitulating epithelial-stromal crosstalk are necessary to understand human reproductive physiology and pathology. Endometrial stromal cells (ESCs) are historically the most utilized primary cell type for experimentation in reproductive biology due to their ability to be cultured and expanded *in vitro*. Recently, protocols to expand and culture primary endometrial epithelial cells as endometrial epithelial organoids (EEOs) have transformed the experimental landscape for *in vitro* analysis of endometrial behavior.^9, 10^ Organoids are self-organizing 3D cell structures that retain many physiologically relevant and functional features of their native tissues of origin.^11^ Thus, EEOs provide opportunities to investigate human reproductive function, understand disease pathogenesis^12^, and test therapeutic compounds for clinical applications.^9, 10^ EEOs are typically generated from isolated primary endometrial glands embedded in Matrigel, a complex, murine tumor-derived ECM primarily comprising heterogenous basement membrane (BM) proteins and growth factors (GFs), together with a defined culture medium designed to enrich the expansion of the stem/progenitor compartment.^13, 14^ Cultured continuously in Matrigel, clonally-generated EEOs retain columnar epithelial architecture of the native tissue and form polarized cyst-like structures that are genomically stable, hormone-responsive, and cell-heterogenous.^9, 10, 15^ However, the reliance on Matrigel to culture organoids presents a critical obstacle for numerous experimental, analytical, and therapeutic applications for dynamic and hormone-sensitive tissues such as the endometrium.^16, 17^ Matrigel’s inherent lot-to-lot variability, poorly-defined molecular composition, limited biophysical properties, and poor suitability for co-culture of epithelia with stromal cells have driven efforts to develop synthetic (or semi-synthetic) ECMs that are fully defined, modular, and tunable.^12, 18–26^

Although endometrial epithelia and stroma are often co-cultured in naturally-derived ECM hydrogels^27–29^, these models are limited by rapid ECM breakdown, which ultimately leads to highly abbreviated experimental conditions and limited control over the biophysical and molecular ECM compositions that can reliably support the morphogenic and functional behavior of multiple cell types. Recently, the development of fully or semi-synthetic ECMs to culture enteric organoids were reported for both murine^23, 30^ and human^17, 21, 26, 30^ tissues. We have previously demonstrated that poly(ethylene glycol) (PEG)-based hydrogel systems can be applied to model some aspects of the endometrial mucosal barrier,^25^ and developed a completely synthetic hydrogel formulation for the culture, differentiation and passaging of human intestinal enteroids that also supported culture of primary endometrial epithelial cells^26^. This synthetic ECM also supports the culture of pancreatic tumor organoids with a complex stroma.^31^ Despite these advances, a fully-defined synthetic ECM that can support the long-term culture and function of multiple endometrial cell populations, especially under the dynamic conditions of hormone variation in a simulated menstrual cycle, is still an unmet need.

Inspired by cell-matrix interactions in native tissue, we designed and synthesized a synthetic ECM that targets cell-specific integrins on epithelia and stroma, supports cell-secreted ECM deposition, and mimics the biophysical properties of the endometrium in health and disease. We show that this bio-labile PEG-based synthetic ECM supports the 3D co-culture of primary human EEOs and ESCs and recapitulates characteristic phenotypic properties of the endometrial responses to sex hormone changes across a simulated 15-day menstrual cycle *in vitro*. We used this 3D co-culture model to parse stromal-epithelial crosstalk and disease-related phenotypes in response to inflammatory cues.

## 2. Methods

### 2.1. Tissue Acquisition and Isolation of Human Endometrial Cells

All participants provided informed consent in accordance with a protocol approved by the Partners Human Research Committee and the Massachusetts Institute of Technology Committee on the Use of Humans as Experimental Subjects (Protocol number IRB-P001994). Endometrial tissue was obtained from pipelle biopsies from reproductive age women (N=18, ages 18-45; Table 1) undergoing laparoscopic surgery for non-malignant gynecologic indications. Study enrollment was limited to pre-menopausal women and excluded patients with an irregular or ambiguous cycle, or a history of hormone use in the 3 months prior to surgery.

Cycle phase dating was performed as described elsewhere^3^ by histological analysis (Fig S20). Endometrial epithelial glands and stromal cells were isolated by enzymatic digestion and filter-separated as previously described^32^, resulting in ≥ 95% purity as assessed by positive staining for vimentin and morphological assessment. Briefly, single stromal cells were separated from intact epithelial glands fragments and cultured separately. Stromal cells were expanded in traditional 2D cultures (as described below) and epithelial glands were cultured as organoids using protocols adapted from Turco et al.^10^ All cells were passaged a maximum of 5 times for all experimental *in vitro* assessments.

### 2.2. Cell Isolation and Endometrial Organoid Generation

Stromal cells were cultured and maintained in phenol red-free DMEM/F12 with 5% charcoal-stripped calf serum, 1 nM 17-β estradiol (E2, Sigma Aldrich) and 1× Pen-Strep solution (Sigma Aldrich). Some cell cultures were treated with 500 nM synthetic progesterone medroxyprogesterone acetate (MPA, Sigma Aldrich). Primary endometrial epithelial organoids (EEOs) were generated from the primary epithelial fragments (p0) according to established protocols and maintained in endometrial epithelial organoid expansion medium (EEO medium).^9, 33^ EEO media is modified from the protocols established by Turco et al.^10^ Briefly, EEO media was defined as the minimal set of components needed to establish organoid cultures expansion and was composed of advanced basal media cocktail^10^ (ABM, 1X); rhEGF (50 ng/mL, Corning); rhNoggin (100 ng/mL, Peprotech); rhRspondin-1 (200 ng/mL, R&D systems); rhFGF-10 (50 ng/mL, Peprotech); Estradiol (E2, 1nM, Sigma Aldrich); Nicotinamide (1mM); Insulin-Transferrin-Selenium (ITS, 1%, Invitrogen); N-Acetyl-L-Cysteine (1.25 mM); TGFB pathway inhibitor (A83-01, 500 nM, Peprotech); and Rock inhibitor Y-27632, (ROCKi, 10 µM, Tocris). For seeding, epithelial cells, as fragments or single cell suspensions, were embedded in 70% Matrigel and cultured for 6 days, with media was changed every two days (Fig S2). For passaging, 6-day old organoids were incubated in Cell Recovery Solution (CRS, Gibco) for 30 mins at 4°C to dissolve Matrigel, then pelleted and digested with Tryp-LE supplemented with 1:100 DNAse (40k units, Sigma) for 15 minutes in a 37°C water bath followed by mechanical dissolution to generate single cells. For expansion purposes, single epithelial cells were seeded in Matrigel at a density of 1,000 cells/µL in 60µL droplets in non-tissue culture treated plates.

We refer to organoids generated from single cells, in regular expansion or in experiments, as scEEOs. For co-culture experiments of ESC and scEEOs, ROCKi was omitted from EEO media since intact organoids were utilized to establish the co-cultures. ESCs retained hormone sensitivity and morphological characteristic when cultured in this EEO-based common media (See Results). Due to the presence of progesterone in the N2 and B27 supplements in the advanced basal media a homemade formulation of EEO media termed neutral EEO (nEEO) media that omitted P4 in the supplements was also explored in some experiments. All media changes for the co-cultures occurred every 3 days for up to 15 days unless noted otherwise.

Spent conditioned media from these cultures was collected and stored at -80°C for downstream analysis.

### 2.3. PEG hydrogel materials and peptides

8-arm poly(ethylene glycol) (PEG) macromers (20 kDa) functionalized with vinyl sulfone (20 kDa PEG-VS) were purchased from JenKem Technology. All peptides were custom synthesized and purified (>95%) by Boston Open Labs (Cambridge, MA), GenScript (Piscataway, NJ), or CPC Scientific (Sunnyvale, CA). Peptides used in these studies include: a dithiol crosslinking peptide containing a matrix metalloproteinase (MMP)-sensitive substrate (Ac)GCRD-LPRTG- GPQGIWGQ-DRCG(Am) (***CL-LW***); fibronectin (FN)-derived peptide containing both the canonical RGD motif from the 10^th^ FN type III domain as well as the PHSRN synergy site from the 9^th^ FN Type III repeat in a branched configuration akin to the biophysical presentation in FN, (Ac)PHSRNGGGK-GGGERCG(Ac) -GGRGDSPY(Am) (***PHSRN-K-RGD***);^34–36^ a Collagen I-derived peptide, (Ac)GGYGGGPG(GPP)5GFOGER(GPP)5GPC(Am) (***GFOGER***);^21, 37^ a peptide with affinity for sequestering cell-produced FN, (Ac)KKGCRE-TLQPVYEYMVGV(Am) (***FN- binder***);^38^ and a peptide with affinity for sequestering the basement membrane proteins type IV collagen and laminin, (Ac)GCRE-ISAFLGIPFAEPPMGPRRFLPPEPKKP(Am) (***BM-binder***).^39^ All peptides were reconstituted in acidic (pH 5.5) Milli-Q water (Millipore). The concentration of free thiols in all peptides was determined using Ellman’s reagent (Sigma Aldrich).

### 2.4. Fabrication of synthetic PEG extracellular matrix

Synthetic matrices were assembled using materials at the concentrations indicated throughout the text. Briefly, 8-arm 20 kDa PEG-VS macromers were prepared in a 10 w/w% solution in ultra pure water. Macromers were used at concentrations ranging from 14.4-33.6 mM VS (nominally, 3-7 wt%). Macromers were functionalized (fPEG-VS) via a Michael-type addition reaction with the integrin (*PHSRN-K-RGS, GFOGER*) and matrix-binding (*FN-binder, BM-binder*) by adding 10% of the total solution volume in 10x PBS + 1M HEPES, pH 8.2, vortex-mixing, and reacting at room temperature for 30 minutes. Note that lyophilized peptides containing HCl salt can be very acidic in stock solutions, so the pH of the final reaction mix should be maintained at 7.8 by adjusting the pH of the buffer solution. During functionalization, cells were counted and pelleted at the appropriate number for the target final cell densities (10,000 ESC and 10 EEO per µL of gel solution). Following functionalization, care was taken to ensure that all cell media was removed without disturbing the cell pellet, and pellets were gently resuspended in fPEG-VS. Upon resuspension, cross linker (CL-LW) was added to the mixture at a stoichiometric ratio of 0.45 thiol:VS to initiate gelation. The solution was gently mixed and plated accordingly, typically in 3 µL droplets in non-tissue culture treated 96-well plates unless otherwise specified in the text. Cell-fPEG-VS mixture was incubated for 25 minutes in a humidified incubator at 37°C until gelation was complete.

### 2.5. Rheological characterization of hydrogels

To measure bulk bio-mechanical properties of crosslinked gels, 25 µL of each gel matrix mixture was loaded into a 1 mL syringe that had the tip cut off at the 0.1 mL mark. The matrix was allowed to gel at 37 °C for 25 min and then extruded from the open syringe as a cylindrical disc and moved to a 24 well plate that contained 400 µL of 1X PBS. The plate was incubated for 24 hours in a humidified incubator at 37 °C, 95% air, and 5% CO2, to allow equilibrium swelling to occur prior to rheological characterization. This procedure generates hydrogel discs of 1.4 mm in thickness. The discs were sandwiched between an 8 mm sandblasted parallel plate geometry and sandblasted base. The shear modulus was determined by performing small-strain oscillatory shear measurements on an Anton Parr MCR 302 rheometer. The mechanical response was recorded by performing frequency sweep measurements (0.1–10 Hz) at a constant strain (0.05), at 37 °C. The elastic modulus (E) is reported as a measure of bulk matrix mechanical properties derived from the recorded storage modulus (G’) in the linear elastic regime where: *E* = 2*G’*(1 + *v*) with the assumption that *v* = 0.5 for ideal elastic hydrogel materials.

### 2.6. EEO generation in synthetic hydrogels

Endometrial organoids were established and expanded in Matrigel according to established protocols^33^ with some minor modifications. Lumenized organoids with a diameter greater than 100 µm were harvested with cold Cell Recovery Solution (CRS, ThermoFisher). Matrigel dollops were manually detached with a wide bore pipette tip, making sure to fully scrape the entire well to lift off the entire gel. The dollops were resuspended in cold CRS (5mL) for at least 30 mins on ice to dissolve remaining Matrigel, followed by centrifugation at 500-1000g, which resulted in a clear pellet of organoids. Organoids were broken into single cell suspensions using 1mL of prewarmed TrypLE Express (ThermoFisher) supplemented with DNAase (10µl, 1:100) and continuous mechanical agitation until a single cell suspension of epithelial cells was observed. Single cells were counted using a hemocytometer then resuspended in the functionalized PEG precursor (fPEG-VS) solutions prior to the addition of the crosslinker. In parallel, single cells were resuspended in Matrigel that served as an experimental control during matrix evaluation. In both cases, cells were encapsulated at a density of 500 cells/µL of matrix. Cell-matrix suspensions were seeded as 3 µL droplets on a non-tissue treated 96 well plate and allowed to polymerize for 25 min in a humidified incubator at 37 °C, 95% air, and 5% CO2. Complete gelation was examined visually on a test gel droplet prior to the addition of media. Gel droplets were maintained in 100 µL of expansion EEO media supplemented with ROCKi (10 µM). Media was changed every three days.

### 2.7. Generation of co-culture model

Six-day old intact organoids (∼100 µm diameter) were grown and expanded in Matrigel and harvested as described above. To maintain scEEO integrity, organoids were incubated in Cell Recovery Solution until Matrigel was fully dissolved. Careful resuspension of intact scEEOs in DMEM/F12 was performed to prevent disrupting scEEO integrity. In parallel, donor matched ESCs were expanded using traditional 2D polystyrene cell culture plates and grown to 80% confluency. ESCs were lifted from 2D culture plates using trypsin (1X) and processed to generate a single cell suspension. Intact scEEOs and ESCs were prepared in parallel to encapsulate both cell types simultaneously and establish a co-culture in the PEG matrix solution. Cells were encapsulated at a density of 10,000 ESC and 10 intact scEEOs per µL of matrix. This epithelial-stromal ratio (approximately 1:1) was determined experimentally based on the cell ratios used to generate established endometrial xenograft models (sub-renal capsule) described elsewhere^40, 41^ and validated using whole-mount 3D image analysis of endometrial tissue we performed (see Results). Embedded cells in fPEG-vs were seeded in 3 µL droplets and allowed to polymerize as described above in non-tissue culture treated 96-well plates. Co-culture was maintained in EEO media without Y-27632, a common media recipe utilized in other co-culture studies^42^, unless otherwise noted. Media was changed every three days and maintained for up to 15 days. It was noted that some basal components of the organoid medium contained P4 (N2 and B27) which may impact hormone responsive behavior, so a P4-reduced ‘neutral’ formulation of this organoid medium (nEEO medium) that completely lacked P4 was tested in some co-cultures. Both media formulations were sufficient to maintain support cell culture and induced hormone-mediated responses (Fig S19); however, as expected, co-cultures maintained in nEEO media conferred a more robust response to progestin. Matching monocultures of ESCs-only or scEEOs-only were established in parallel for some experiments. Cultures were imaged daily.

### 2.8. EEO and co-culture efficiency and quantification

To quantify organoid diameter and total organoid formation efficiency, single epithelial cells were encapsulated in fPEG-VS hydrogels or Matrigel and cultured over 14 days. scEEO formation efficiency was calculated as the percentage of organoids with a clear lumen with a diameter (>100 um) relative to Matrigel. 4X brightfield (BF) z-stack images of 10-day old endometrial organoids were captured using an EVOS M500 microscope (Invitrogen). The images were processed in Fiji using the time lapse Gaussian-based stacker focuser plugin to generate maximum intensity projections. Images were manually analyzed in Fiji to obtain a diameter and organoid count distribution for each condition. Daily images of co-cultures were manually curated to obtain relative growth rates of the organoids in single culture (N=3) or co-culture (N=8) by measuring scEEO diameter and quantifying the daily fold change relative to day 1 of culture (video 3, 4).

### 2.9. Cell viability and proliferation analysis

Co-culture cell viability and cell death was measured using the LIVE/DEAD™ Cell Imaging Kit (488/570) (Thermofisher Scientific) according to the manufactorers protocol. Briefly, At day-15 prior to fixation, cells were incubated the reagents provided by the kit and incubated at 37 °C, 95% air, and 5% CO2 for 20 minutes. Images were captured using Keyence BZ-X800 Fluorescence Microscope. Cell proliferation was assessed by EdU incorporation using the Click-iT Plus EdU imaging kit with Alexa Fluor 594. All co-culture and monoculture experiments were treated with EdU (5-ethynyl-2′-deoxyuridine, 20 μM) for a 24-hour incubation period at 37°C, 95% air, 5% CO2 prior to fixation at day 15 of culture. Click-iT reaction cocktail was prepared as described by the manufacturer (ThermoFisher Scientific). Cells were incubated for 1 hr at RT protected from light and counter stained with DAPI (1 mg/mL, 1:2000). Images were captured using a ZEISS confocal Laser Scanning Microscope (LSM 880) and analyzed by 3D image processing was performed using the Surface and Spots function in Imaris (Bitmap) software (9.6.0). At least three individual organoids were analyzed per donor, and the proliferative index was measured as the number of EdU^+^ positive cells per organoid surface area (mm^2^). ESC proliferative index was calculated as the EdU^+^ cells relative to the total number of DAPI stained nuclei.

### 2.10. RNA isolation for quantitative real time PCR (qPCR) and bulk RNA-seq

ESCs and intact scEEOs cultured in Matrigel or synthetic hydrogels were used for molecular analysis. After 14 days of culture, intact scsEEOs were released from Matrigel and the synthetic ECM, using Cell Recovery Solution, or recombinant Sortase, respectively. Intact scEEO-only cultures, ESC-only cultures, or co-cultures were pelleted, resuspended in TRIzol reagent (ThermoFisher Scientific), and then stored at -80°C until processing. RNA was extracted using the Directzol RNA Mini-Prep kit (Zymo Research) per the manufacturer’s protocols with the inclusion of an on-column DNase step using the PureLink DNase Set (ThermoFisher Scientific). At least eight replicates (3 µl droplets) were pooled together to obtain sufficient RNA. cDNA was synthesized from ∼1 µg of total RNA using the High-Capacity RNA-to-cDNA Kit (ThermoFisher Scientific, 4387406) per manufacturer’s protocols. TaqMan Fast Advanced Master Mix (ThermoFisher Scientific) was used in congruence with the cell-specific probes for qPCR. Gene expression was determined using the StepOnePlus real-time PCR system (Applied Biosystems) and calculated using the ΔΔCt method using GraphPad Prism. Gene expression was first normalized using the housekeeping GAPDH gene in each sample, then the relative fold change relative to experimental control. High Throughput 3’ Digital Gene Expression (3’DGE) RNAseq was performed on the isolated RNA using NovaSeq flowcells at the BioMicroCenter (MIT) and processed using the BMC/BCC 1.8 pipeline (cocultures n=7; ESC monocultures, n=6).

### 2.11. Secreted protein measurement by ELISA and Luminex

Spent media was stored at −80°C until analyzed by ELISA for prolactin (R&D Systems). Protocols provided by the manufacturer were adapted to allow ELISAs to be performed in a 384-well plate (ThermoFisher) to minimize medium (sample) volume needed.^25^ Multiplexed Luminex assays were performed to measure cytokines in 48- or 72-hour undiluted conditioned medium throughout the co-culture period using the magnetic Human Luminex XL Cytokine 45- PLEX pre-mixed kit (R&D systems). Protocols provided by the manufacturer were adapted to allow the assay to be performed in a 384-well plate to avoid introducing batch variability. Ten-point standard curves with culture medium and assay buffer blanks were included for quantification. Conversion from mean fluorescence intensity (MFI) values to absolute concentration was performed by interpolating 5-parameter logistical fits of standard curves for each analyte. Following the same downstream analysis protocol, secreted matrix metalloproteinases (MMP) concentrations were measured using the magnetic Human MMP Magnetic Luminex Performance Assay (R&D Systems). This assay required dilutions of cell culture supernatants in fresh culture medium at either 2x or 200x to detect the reported analytes within a quantifiable range.

### 2.12. Immunostaining and image analysis

Hydrogel samples were fixed in 4% paraformaldehyde (Electron Microscopy Sciences) for 30 mins, permeabilized in 0.2% TritonX-100 in PBS (Sigma) for at least 30 mins at room temperature, and then blocked overnight with blocking buffer (1% BSA (Sigma), 5% normal donkey serum (Electron Microscopy Sciences) in PBS) at 4°C. Hydrogels were incubated with primary antibodies overnight while rocking at 4°C, rinsed with blocking buffer 3 times (5 min each while rocking at RT), and treated with secondary-conjugated antibodies overnight while rocking at 4°C. Specific antibodies can be found in the supplementary table of reagents. Images were captured using a ZEISS confocal Laser Scanning Microscope (LSM 880) or a Keyence BZ-X800 Fluorescence Microscope. Image analysis was performed using Fiji open software and 3D image processing was performed using Imaris (Bitmap) software (9.6.0).

### 2.13. Endometrial ECM proteomics

Endometrial ECM processing for proteomics, proteomics data collection, processing and analysis is described elsewhere.^43^ Briefly, fresh endometrial biopsies were collected and processed using 50 µl of RapiGest (Waters Corporation) reconstituted in 50 mM NH4HCO3 at a concentration of 2mg/ml to digest the endometrial biopsy sample for 24 hours at 37°C. Next, urea (ThermoFisher) and dithiothreitol (ThermoFisher) were added to the sample to bring the mixture of a final concentration of 4 mM and 5 mM, respectively. Finally, iodoacetamide (MilliporeSigma) was added to the sample, for a final concentration of 15 mM and kept in the dark, at 37°C, for 30 minutes. Samples were frozen at -80°C until all patient samples had been collected and processed. Samples were lyophilized overnight and then reconstituted in NH4HCO3. Samples were digested with Trypsin Gold, mass spectrometry grade (Promega), using a ratio of 1 µg trypsin to 50 µg sample protein as quantified using a Pierce BCA Protein Assay Kit (ThermoFisher). Samples were digested overnight at room temperature using a rotor for mixing and then placed in a speed vacuum to reduce sample volume. Dried down digested peptide samples were labeled with TMT10plex kit (Pierce). After labeling, the samples were mixed and purified with peptide desalting spin columns (Pierce). 1/5 of the sample was used for one LC/MS/MS analysis (Thermo Exploris480 mass spectrometer). The dried peptide mix was reconstituted in a solution of 2% formic acid (FA) for MS analysis. Peptides were loaded with the autosampler directly onto a 50cm EASY-Spray C18 column (Thermo Scientific). All MS/MS samples were analyzed using Sequest (Thermo Fisher Scientific; version IseNode in Proteome Discoverer 2.3.0.523). Sequest was set up to search uniprot_human_reviewed_032120.fasta (version March 21, 2020) assuming the digestion enzyme trypsin. Scaffold Q+ (version Scaffold_5.0.1, Proteome Software Inc.) was used to quantitate Label Based Quantitation peptide and protein identifications. Peptide identifications were accepted if they could be established at greater than 10.0% probability to achieve an FDR less than 1.0% by the Percolator posterior error probability calculation.^44^ Normalization was performed iteratively (across samples and spectra) on intensities, as described in Statistical Analysis of Relative Labeled Mass Spectrometry Data from Complex Samples Using ANOVA.^45^ Medians were used for averaging. The protein list was then compared to the curated human matrisome (http://matrisome.org/) list to look for ECM proteins. Samples were normalized using a mixed aliquot of all patient samples that was analyzed in each LC/MS-MS run. Overlapping ECM proteins found in all 3 LC/MS-MS runs were identified and the Limma package^46^ in R was used to remove batch to batch effects.

### 2.14. Endometrial tissue single cell RNA sequencing (scRNAseq)

scRNAseq data collection, processing and analysis as described elsewhere (in preparation)^43^. Data was downloaded and Seurat (V3) was used to create dot plots and violin plots of specific genes of interest on scaled data.

### 2.15. Bulk RNAseq analysis of in vitro co-cultures

Aligned RNAseq transcript expected counts were analyzed in R (version 4.0.0) where data was modeled using the edgeR package (version 3.34.0).^47, 48^ Counts were normalized by library size and filtered to exclude genes with less than 20 transcripts across all samples. Principal component analysis was performed using data batch-corrected by donor via the ComBat-seq method of the sva package (version 3.13) in R, and each gene was scaled to have unit variance before modeling.^49^ Differential gene expression was calculated by modeling transcript data in a quasi-likelihood negative binomial generalized log-linear model using the edgeR function “glmQLFit”. The use of a generalized log-linear model allowed for differences between hormone treatment groups to be compared while accounting for inter-donor variability. Genes were ranked by fold change in expression between hormone treatment groups to produce a ranked list of genes for gene set enrichment analysis (GSEA) (http://www.broadinstitute.org/gsea/index.jsp). Gene ontology (GO) term analysis was performed on ranked gene lists of genes with pvalues <= 0.05. No adjustment was performed at this step due to the low sample number, high variability, and intent to use the RNA sequencing as a hypothesis generation step. The R package topGO (version 2.44.0) was used for GO term identification and the package clusterProfiler (version 4.0.2) was used for visualizations. A curated list of menstrual related genes was assembled from existing literature.^50, 51^ A heatmap of log_"_ transformed data was created using the R package pheatmap (https://CRAN.R-project.org/package=pheatmap; version 1.0.12) with the “complete” clustering method. Analysis was performed on 7 co-culture experiments comprised of 5 donors, 2 independent technical replicates and results were compared between experiments that were maintained in nEEO media (N=2) or traditional EEO media (N=5).

### 2.16. Statistical analysis

Data are expressed as average ± standard error of the mean (SEM). Statistical analysis was performed using GraphPad Prism 8. Unpaired t-tests assuming unequal variances and two-way ANOVAs with appropriate post-tests were performed. Holm-Sidak’s multiple comparison of the mean or two-tailed Mann- Whitney tests when comparing two groups. qPCR data was analyzed using multiple t-tests. Statistical significance was defined as * p < 0.05, ** p < 0.01, *** p < 0.0001, and ****P < 0.0001.

## 3. Results

### 3.1 Design of a fully synthetic ECM guided by multi-omic evaluation of the human endometrium

The ECM composition of the endometrium, along with cell-matrix receptor expression, are regulated by sex steroids throughout the menstrual cycle,^52, 53^ giving rise to dramatic shifts in tissue structure and morphology (Fig 1A). The overarching design goal for our synthetic ECM is to define a minimal set of biophysical and biomolecular cues required to support the dynamic phenotypic functions of heterogenous endometrial cell populations, over weeks of culture.

As a first step, we performed a targeted proteomic analysis (LC-MS/MS, N=11) of the endometrial ECM-associated proteins (matrisome) across the menstrual cycle to benchmark the ECM composition in the native tissue (Fig 1B). Consistent with immunohistochemical reports^54–,56^, FN abundance was greater in the proliferative phase, while fibrillin, collagen and laminin proteins increased during the secretory phase, corresponding to the differentiation of stromal fibroblasts, epithelial gland maturation, and vascular remodeling in response to progesterone (Fig 1B).

Our initial studies demonstrating the establishment of endometrial organoids in a synthetic ECM^26^ were guided in part by published histologic analysis of endometrial integrin expression patterns from a subset of the 24 known integrins.^37, 56–62^ We sought additional insights into cell population-specific integrin expression by probing a single cell RNA-sequencing dataset^43^ (scRNAseq, N=6; Fig 1C; Fig S1) to profile the integrin expression across the endometrial proliferative or secretory phase epithelia and stroma in addition to the immune and vascular cellular compartments. Across all cells in the data set, transcripts for all eighteen α chains and all eight β chains, except integrin β3, were detected (data not shown). Integrin β3, a contested marker of fertility that appears in the secretory phase^63–65^, was also absent in another endometrial scRNAseq data set.^66^ Consistent with the immunohistochemistry literature^58^, αV was robustly expressed by epithelia and stroma mostly during the proliferative phase. Its primary binding partner β5 was primarily expressed by the stroma across the cycle (Fig S1A), while β6 was sparse except in proliferative phase epithelia (Fig S1A). Unexpectedly, integrin β8, an additional αV partner that has not previously been characterized via immunohistochemistry in the endometrium, was robustly expressed in several epithelial and some stromal subpopulations (Fig 1C; Fig S1A). Also consistent with the immunohistochemistry data^56, 59–61, 67^ integrin β1 was robustly expressed by both epithelia and stroma across the cycle. Expression of the collagen-binding α1 dimer (at lower levels, especially in proliferative epithelia) along with strong expression of α2 in the epithelia was also observed, while the stroma primarily expressed α5 (Fig 1C, Fig S1B). Finally, laminin-binding α6 and its heterodimer β1 was robustly expressed in all epithelia, while expression in the stroma was higher in the proliferative phase (Fig S1). We investigated the expression of a subset of these integrins in cultured endometrial cells via qPCR confirming comparable expression of β1, α1, α2, α8 and αv in both stromal and epithelial cells, and significantly greater expression of α5 and α8 in stromal cells (Fig S1C).

The minimal synthetic ECM must provide integrin-engaging adhesions cues needed by the cells to support viability immediately post-encapsulation upon dissociation from tissue or a prior culture state^68, 69^. We thus include the peptide *GFOGER*, recognized by α1β1 (expressed on stroma and epithelia) and α2β1 integrin heterodimers (expressed on epithelia),^70–72, 73^ together with the fibronectin-derived *PHSRN-K-RGD* peptide ^4, 26^ as an RGD ligand for αVβ1 and α5β1 (Fig 1D). This ligand incorporates the PHSRN synergy site recognized by α5β1 heterodimer primarily expressed by the stroma and, to a lesser extent, by the secretory epithelium (Fig 1C).

As cells remodel the local microenvironment, proteolytically degrading the synthetic ECM, they deposit cell-specific ECM, creating a dominant source of adhesion cues for a broad spectrum of additional adhesion receptors as the culture progresses. To sequester this cell-produced ECM and enhance its interactions with the synthetic ECM, we incorporated peptides that bind to basement membrane components produced by epithelia (and some decidual fibroblasts) and to fibronectin produced by stromal cells, as previously described.^25, 26, 31^ This strategy obviates the need to include laminin-derived peptides.^25, 26, 31^

To synthesize the synthetic ECM, we first functionalized an 8-arm poly(ethylene glycol) (PEG)-based hydrogel with a combination of *GFOGER* and *PHSRN-K-RGD* (“*MIX”*) in defined ratios (see below), then crosslinked this macromer solution, in the presence of cells, with a protease-degradable crosslinking peptide (“*CL-LW”*) to enable cell-dependent matrix remodeling, migration and proliferation during hormone-driven morphogenesis.^25, 35, 74^ Using this basic framework, individual synthetic ECM parameters (e.g., matrix stiffness) were systematically and independently varied to mimic aspects of healthy and disease states (Fig 1D). Altogether, a tissue-inspired approach was implemented to design a synthetic matrix that would initiate and maintain the culture of EEOs and ESC in a controlled environment that mimics the molecular and biophysical properties of the endometrial ECM.

### 3.2 Synthetic ECM supports 3D co-culture of primary human endometrial stromal and epithelial cells

#### 3.2.1. A semi-empirical screen of synthetic ECM properties yields a cue-response phenotypic landscape for endometrial epithelial organoids

We previously demonstrated that a particular formulation of synthetic ECM designed for enteric organoids also supported endometrial epithelial organoids^26^; however, the impact of the biophysical properties and peptide composition on organoid phenotype and function was largely unexplored. These first-generation hydrogels^26^ incorporated only one adhesion ligand (i.e., *GFOGER*) and were substantially stiffer (∼2000 Pa, corresponding to 5wt% PEG gels) than Matrigel (∼150-443 Pa)^20, 75^ or the native endometrial tissue (∼250 Pa)^76^. Thus, to better understand EEO generation in synthetic matrices, we probed the biophysical parameters that more closely mimicked native endometrial stiffness and additionally interrogated the consequences of integrin ligand composition. We first generated a tissue bank of EEOs from 8 endometrial donors, with and without endometriotic disorders (Table 1; Fig S2A) and defined a panel of gels with varying elastic moduli, ranging from “soft” gels (3wt% PEG, ∼ 300 Pa) comparable to native stiffness of the endometrium, to the “stiffest” gels (7wt% PEG, ∼6,000 Pa; Fig 2A). To test these formulations, we dissociated organoids to single cells, passaged them into synthetic matrix or Matrigel, and followed the emergence into single cell-derived endometrial epithelial organoids (“scEEO”, Fig S2B). In agreement with our previous work^26^, employing synthetic hydrogels functionalized with *GFOGER* was sufficient to support the growth of scEEOs, even in a relatively stiff (5wt% PEG) environment. However, scEEOs generated in the stiffer gels developed a crenelated morphology (Fig 2B; Fig S4) compared to those cultured in softer conditions (3wt% PEG), and this was consistent across all donors regardless of the donor’s disease state (data not shown).

**Figure 2.**
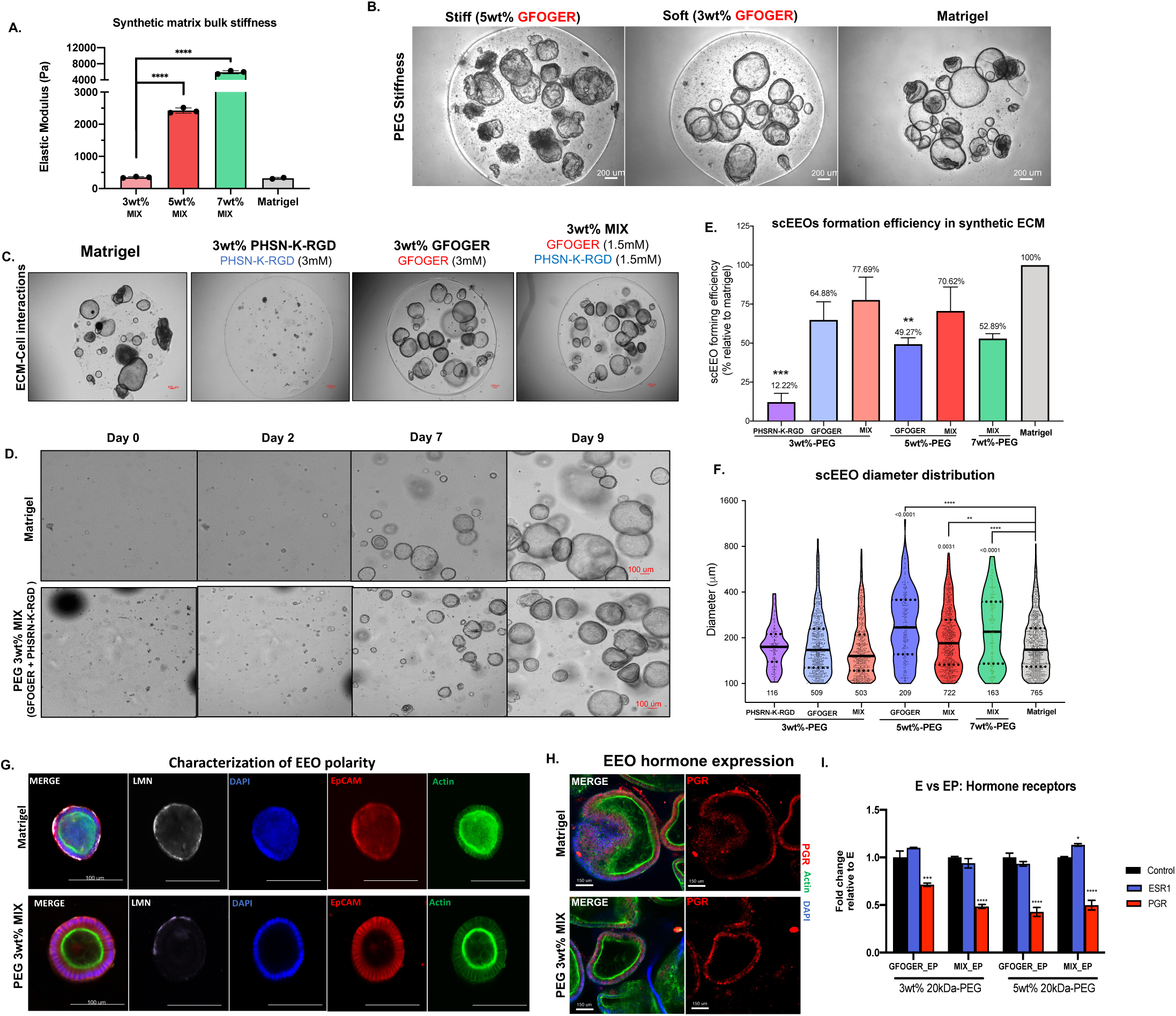
Dual integrin ligand functionalization in softer synthetic matrix regimes enhance EEOs generation. (A) Biomechanical properties (elastic moduli, Pa, n=3 gels) of synthetic hydrogel formulations, where “soft” = 3wt% (∼300 Pa), “stiff” = 5wt% (∼2kPa) and “stiffest” = 7wt% (∼6kPa). (B) Representative images of scEEO and morphology in stiff (5wt%) in soft (3wt%) PEG hydrogels functionalized with only GFOGER (3mM) compared to Matrigel. (C) Day-15 images of hydrogel formulations comparing adhesion peptides reveals that GFOGER is necessary to generate scEEOs. Dual-adhesion-peptide functionalization with GFOGER (1.5mM) and PHSN- K-RGD (1.5 mM) is sufficient for robust scEEO generation. (D) Time-lapse images of 10-day cultures of EEO derived from single cells in synthetic hydrogels (PEG 3wt%-MIX). (E) Quantification of the scEEO formation efficiency in synthetic hydrogels relative to Matrigel (N=8). (F) Quantification of the total lumenized EEO (>100 um diameter) count and EEO diameter distribution after 10 days in culture. (G) Characterization of EEO morphology and polarity of EEOs generated in synthetic ECM (3wt%-MIX) compared to Matrigel via immunofluorescent (IF) analysis of ECM deposition for laminin (LMN) and F-actin localization after 14 days of culture in soft (3%) PEG MIX hydrogel and in Matrigel. (H-I) Characterization of protein and transcriptomic expression of hormone receptors in EEOs generated in synthetic hydrogels. (H) Representative images of PGR staining countered stained with F-actin (green) and DAPI (nuclei) and (I) qPCR analysis of mRNA expression of PGR and ESR in EEOs treated with either E2 (E) or E2 + MPA (EP) for 14 days. Analysis compares ESR, PGR values relative to matching E treatment groups (control). Significance is indicated as *p<0.05, **p<0.01, ***p<0.001.

Anticipating the integration of α5β1-expressing stromal cells, we then evaluated the influence of an additional integrin ligand on scEEO growth behavior and morphologies. We compared ECM formulations incorporating nominal concentrations of *GFOGER* (3mM), *PHSRN-K-RGD* (3mM), or a *MIX* of both peptide (1.5 mM each) in the soft (∼300 Pa, 3wt% PEG) gel regime. Biomechanical properties were not significantly affected by differences in integrin ligand composition (data not shown). As expected, *GFOGER* was necessary to generate scEEOs in synthetic matrices (Fig 2C; Fig S3), presumably through engagement of α1β1 and α2β1, and although *PHSRN-K-RGD* conferred minimal scEEO generation alone, incorporation of both adhesion ligands in the *MIX* formulation robustly promoted the growth of the scEEO despite the lower (1.5 mM vs 3 mM) concentration of GFOGER in the *MIX* (1.5 mM *GFOGER* + 1.5 mM *PHSRN-K-RGD*) formulation. When visualized by brightfield imaging on sequential days after seeding single cells, organoids appeared to emerge more slowly in PEG matrices (soft, 3wt%) than in Matrigel, yet they retained similar morphologies and diameters by day 9 of culture (Fig 2D) suggesting that the 3wt% gels function with comparable efficacy to Matrigel.

Having confirmed the requirement for *GFOGER*, and comparable scEEO emergence in both *GFOGER*-only and *MIX* gels, we next compared how matrix stiffnesses impacted scEEO emergence by varying the PEG polymer content from 3wt% (∼300 Pa) to 7wt% (∼6 kPa) to mimic physiologic^76^ and pathologic^77^ uterine regimes, respectively (Fig 2E-F). The efficiency of scEEO formation, as assessed by the number of organoids at day 14 relative to Matrigel, was slightly inversely correlated with matrix stiffness (Fig 2E). Moreover, organoid diameter distributions in stiffer matrices, where organoids were sparser, skewed slightly higher in the 5wt% and 7wt% gels compared to softer 3wt% matrices (Fig 2F). Consistently, the organoids cultured in stiffer matrices (5wt% and 7wt%) manifested the meandering, crenelated morphology described above, but not in softer PEG matrices or Matrigel (Fig S4). These results suggest that EEO clonal establishment is robust across a range of biophysical properties (Fig 2E-F) and provide a preliminary indication that tissue stiffness influences epithelial morphology and behaviors.

Based on these results, we focused further investigations using the more physiologically relevant 3wt%-*MIX* gels to characterize scEEO architecture, hormone response and cellular heterogeneity in the synthetic ECM. PEG-derived scEEOs exhibited proper polarity based on basolateral laminin (LMN) deposition and apical F-actin accumulation, comparable to those generated in Matrigel (Fig 2G). Moreover, scEEOs generated in synthetic matrices retained their epithelial origin and architecture as assessed by EpCAM and F-actin staining (Fig S5A-B), were mitotically active (Fig S5C), and exhibited appropriate response to sex steroids as demonstrated by immunohistochemical staining of progesterone receptors (PGR) (Fig 2H) and the downregulation of PGR transcript, but not ESR, in response to progestin treatment (Fig 2I). To confirm scEEO maturation, live-imaging and immunostaining revealed distinct populations of motile ciliated and secretory epithelial cells (Fig S5D, Video 1). These results were consistent across several tissue donors (N=8). Altogether, these results justified the use of soft (3wt%) PEG hydrogels, functionalized with a dual presentation of *GFOGER* and *PHSRN-K-RGD* (*MIX*), as a functional synthetic ECM to generate and maintain long-term cultures of endometrial organoids (Fig 1D).

#### 3.2.2. Establishment and characterization of ESC 3D cultures in synthetic ECM

Our results examining organoids in synthetic matrices demonstrated that both the biophysical and the molecular parameter space of the hydrogels are important to promote adequate scEEO growth and function; therefore, we set out to evaluate, in parallel, how adhesive cues and matrix stiffness influenced ESC behavior in 3D (Fig 3A). Embedded ESCs cultured in the gels were viable in all conditions tested (*GFOGER*, *PHSRN-K-RGD*, or *MIX* for the 3-7% PEG) and maintained comparable cell numbers across all conditions (data not shown). However, ESCs adopted a more characteristic stellate morphology in softer (3wt%) hydrogel microenvironments compared to the stiffer (5wt%) gels which constrained fibroblast spreading and outgrowth (Fig 3B; 7% not shown). Furthermore, in support of previous studies^25^, hydrogels containing *PHSRN-K-RGD* further provided a modest increase in stromal dispersion and spreading, as assessed by the fluorescence projected area covered on day 7 of culture, compared to matrices containing *GFOGER* alone where cells retained a more clumped distribution (Fig 3C).

**Figure 3.**
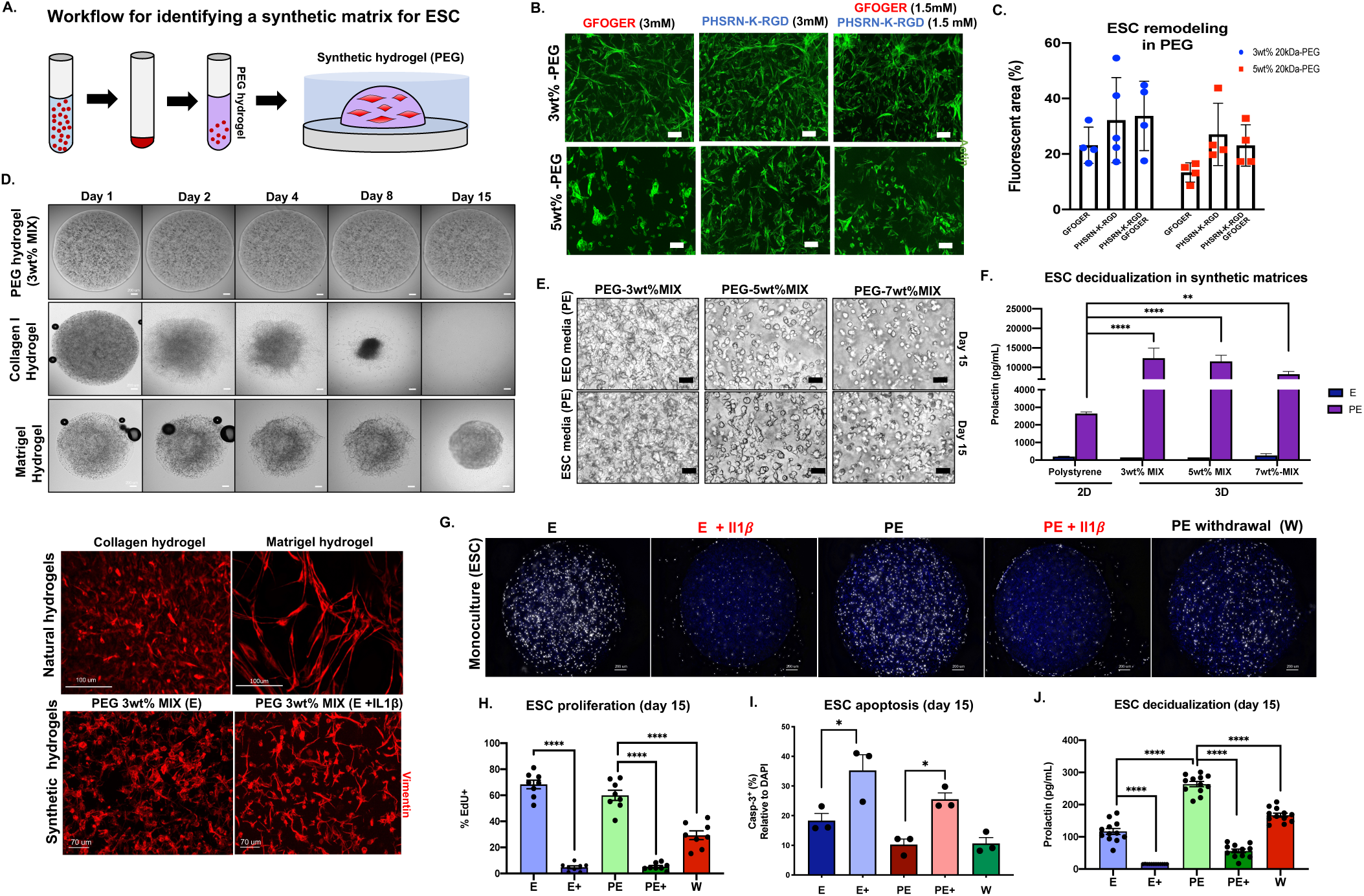
Development and characterization of a synthetic endometrial stromal cell culture. (A) Schematic of the workflow for culturing ESCs in 3D using a synthetic hydrogel. (B) Maximum intensity projection images of F-actin (green) reveal ESC remodeling (cell elongation and dispersion) at day 7 of culture in stromal (serum containg) media impacted primarily by hydrogel stiffness (20X magnification) as quantified in (C). (D) Time-lapse images of embedded ESCs in synthetic matrix (PEG) compared to the natural-derivedhydrogels Collagen I (∼4mg/mL) and Matrigel (8 mg/mL) allow for long-term stable cultures. Vimentin stain (red) of ESC morphology cultured in synthetic and natural hydrogels. IL-1β (1ng/mL) treated ESC in synthetic hydrogel provided as a control for inflamed morphology. (E) Brightfield images of stromal decidualization morphology in synthetic ECMs as a function of matrix stiffness (3wt%, 5wt% and 7wt% *MIX*) after 14 days of EP treatment and in standard ESC media or EEO medium (see Methods). (F) Biochemical measurement of prolactin (PRL) in the spent media (EEO media) from 15-day cultures in (E) compared to standard 2D polystyrene cultures (N=4). (G) Represenatitive fluorescent staining of DNA synthesis (EdU incorporation for 24 hrs hrs prior to fixation at day 15 as a function of simulated cycle phase and inflammation cue, IL-1β (H) Quantification of mean EdU^+^ staining from cultures shown in (G) shows IL-1β suppresses DNA synthesis of ESCs (n=3) (I) Quantification of apoptosis (cleaved caspase-3^+^) in ESC cells stimulated with homrone and IL- 1β treatment groups at day 15 of culture relative to total nuclei count (N=3) (J) Inflammatory challenge with IL-1β significantly suppressed PRL secretion in ESC cultures (E+ = E + IL1β, PE+= PE+IL1β, W = hormone withdrawal) (N=6). Significance is indicated as *p<0.05, **p<0.01, ***p<0.001. Scale bars are 50 μm, unless otherwise noted.

Although the ESCs can enzymatically remodel the synthetic matrices due to the incorporation of the MMP cleavable crosslinker, PEG gels remained intact at the 15-day time point, whereas cultures in Matrigel or collagen I gels have shrunken and at least partly disintegrated by this time point (Fig 3D). This ability to maintain long-term cultures enabled us to assess stromal function in the gels by maintaining the 3D cultures under sex hormone stimulation throughout the 15-day experiment. In response to treatment with E2 (E), a synthetic progestin (E2 + MPA, “PE”), or a PE hormone withdrawal (72 hrs, ‘W’), subsets of ESCs underwent decidualization, a differentiation process necessary for the establishment of pregnancy, characterized by the secretion of pro-gestational protein prolactin (PRL) and morphological changes toward an epithelial-like decidual cell.^42, 51, 78^ Although increased matrix stiffness curtailed ESC elongation in the E-containing medium before decidualization, ESC developed characteristic rounded epithelioid morphologies in response to progestin in the *MIX* gels, in both standard ESC culture media as well as EEO media (Fig 3E). Detection of PRL secretion in the spent media confirmed that ESC embedded in 3D gel underwent robust decidualization in all matrix stiffnesses response to progestin treatment (Fig 3F), suggesting that stiffness is not a strong influence on this metric of stromal hormone sensitivity.

We next investigated the effects of exogenous treatment with IL-1β (1 ng/mL), a pro-inflammatory cytokine associated in reproductive disorders^79^, on ESC phenotype in the synthetic gels. Compared to control groups, which exhibited robust proliferation at day 15 in the 3wt% *MIX* hydrogel in both E and PE media, as 50-60% of cells incorporated EdU over a 24- hour incubation period (Fig 3G-H), IL-1β dramatically suppressed ESC mitotic activity resulting in ∼5% of cells incorporating EdU (Fig 3G) and induced an increase in apoptosis (Fig 3I). Furthermore, by day 15, treatment with IL-1β also dramatically suppressed PRL production even in the presence of the progestin MPA (Fig 3J). Brightfield and immunofluorescent examination of stromal morphology in response to the inflammatory cue revealed a drastic change toward an elongated morphology, characterized by a thin architecture (Fig S6A-B) and decreased cell surface area (Fig S6C). This change was attributed to an apparent increase in ECM remodeling capacity characterized by increased cellular motility (Video 3). These results suggest that despite a reduction of mitotic activity, IL-1β induces a more activated ESC phenotype in 3D. In line with this result, morphological assessment revealed that ESC cultured in Matrigel also developed the phenotype akin to those exogenously treated with IL-1β (Fig 3D), highlighting the limitations conferred by Matrigel to study stromal behavior^27^. Altogether, these results demonstrate that the relatively soft, dual integrin ligand (3wt% *MIX*) synthetic matrix formulation is suitable as a “one size fits all” scaffold for culturing both stromal cells and epithelial organoids.

#### 3.2.3. Engineered synthetic matrices support the stable co-culture of EEO and ESC, capturing the temporal physiologic processes of the human menstrual cycle

In the endometrium, both stromal and epithelial cells express hormone receptors for estrogen and progesterone, yet proper tissue-level function is mediated by the crosstalk between these cell types.^80–83^ Parsing this crosstalk *in vitro* using Matrigel - which contains numerous exogenous growth factors, rapidly degrades, and largely lacks the physiologic ECM molecules for stromal cells – is arguably a fraught endeavor. Thus, we used the fully defined synthetic matrices described to develop a co-culture model using defined cell ratios that were shown to approximate those observed *in vivo* (Fig S7) to interrogate these dynamic phenomena. Donor matched EEOs and ESCs populations (N=12) were co-cultured in the synthetic ECM (3wt%-*MIX*) using a fully defined common media (see Methods) and maintained for up to 15 days of culture (Fig 4A; Video 2,3). Immunostaining demonstrated the persistence of morphologically well-defined stromal and epithelial populations (Fig 4B; Video 2). All cultures were viable, with minimal cell death, throughout the length of the experiments as demonstrated by live/dead staining (Fig S12) at day 15. Co-cultures were exposed to hormone stimulation designed to mimic the proliferative (E2, ‘E’), secretory (E2 + MPA, ‘PE’), or menstrual (72 hr PE withdrawal + RU-486, ‘W’) phases of the idealized human menstrual cycle (Fig 4C-D).

**Figure 4.**
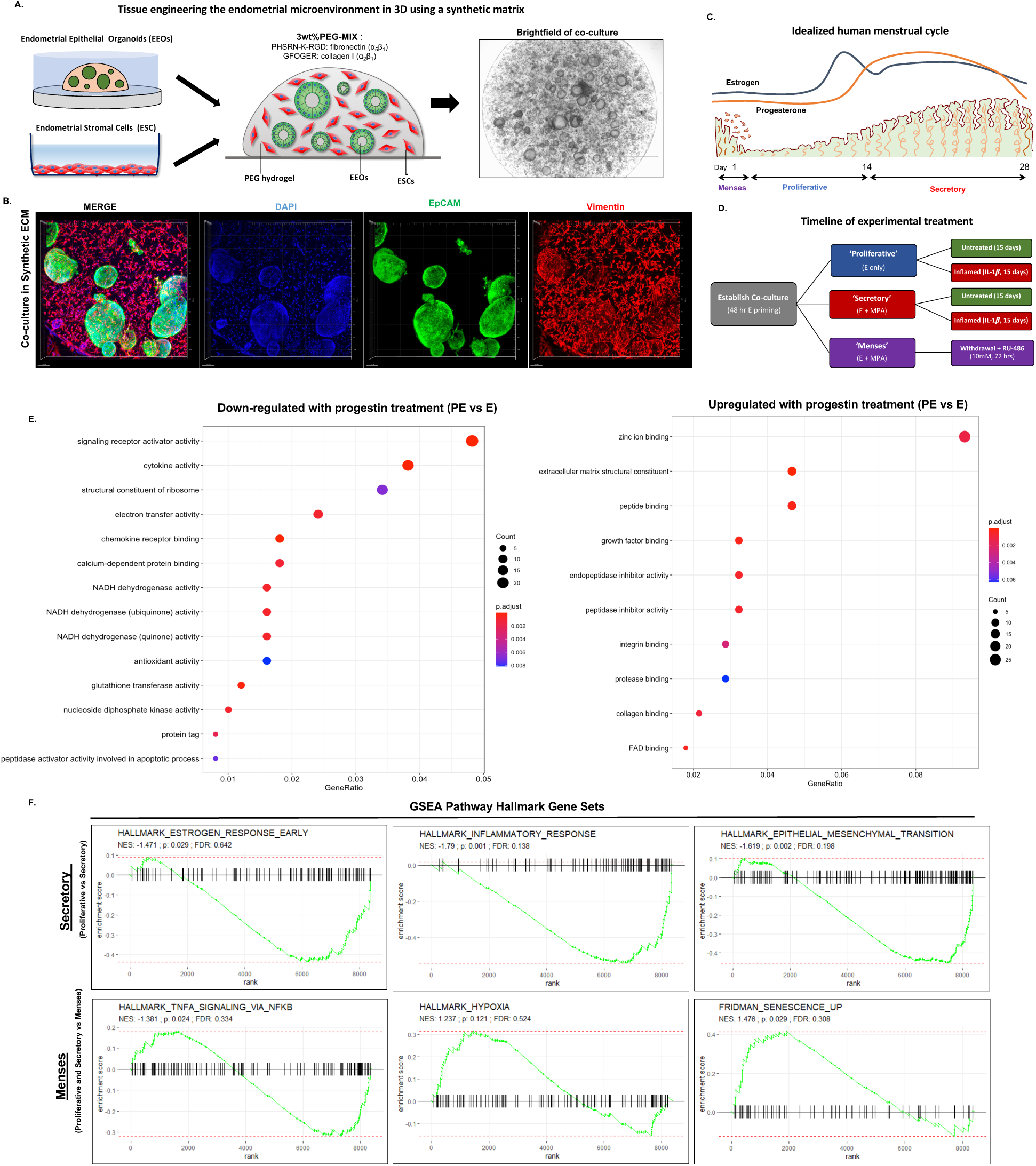
Establishment of a co-culture model recapitulates the molecular signature of the human menstrual cycle in vitro. (A) Schematic and representative images of the co-cultures. EEOs (10 intact organoids/uL) and ESC (10k cells/uL) are expanded separately and embedded in the synthetic ECM, cultured as 3ul droplets and maintained for 15 days of culture. A dual- adhesion peptide synthetic ECM (3wt% 20kDa-PEG) functionalized with GFOGER (1.5mM) and PHSRN-K-RGD (1.5mM) supports the establishment of the EEOs and ESC co-culture model. (B) Representative 3D images of day-15 endometrial EEOs (EpCAM = green) and ESC (Vimentin = red) in the co-cultures (scale =100 um). (C) Schematic and timeframe of the idealized 28-day menstrual cycle in vivo. (D) Experimental design for the validation of the co-culture endometrial model. Treatment groups are designed to mimic the hormonal changes in the phases of the menstrual cycle, including a pharmacologic induction of the menstruation using a PGR antagonist (RU-486, 10uM). ‘Inflamed’ groups were co-treated with IL1β (1ng/mL) throughout the length of the experiment. (E-G) Transcriptomic (bulk RNAseq) analysis of the co-culture models (n= 7). (E) Gene ontology (GO) analysis of the top significantly enriched GO terms that are downregulated or upregulated in response to progestin treatment. Cocultures treated with E2 + MPA (“PE”) have gene ontology (GO) expression profiles that align with the secretory phase of the menstrual cycle. GSEA analysis of hallmark pathway gene sets show pathway changes consistent with corresponding phases of the menstrual cycle. All images and transcriptomic analysis were performed on day-15 co-cultures in the synthetic 3wt% 20kDa-PEG-Mix hydrogels (N=8). Data is shown with the −log of their P values.

To determine whether the PEG-based co-culture model could phenocopy canonical molecular pathways involved in the human menstrual cycle, we characterized the co-cultures (N = 5 donors) by both transcriptomic and functional analysis after hormonal treatment. First, we implemented bulk RNA sequencing (n=7) and demonstrated that after 15 days, co-cultures were transcriptionally distinct from either ESC or EEO monocultures (Fig S8B, Fig S19). Further analysis using a generalized linear model compared gene expression between ESC monocultures and co-cultures using Benjamini-Hochberg (BH) multiple hypothesis test correcting^84^, resulted in 7,821 genes to be differentially expressed at an adjusted p-value threshold of 1×10^-5^ (Fig S8C). These findings are suggestive of a crosstalk between these stromal and epithelial populations when cultured together.

Next, we analyzed the response of co-cultures with respect to hormonal signaling pathways. Further transcriptomic analysis of the co-cultures by hierarchical clustering demonstrated that the expression of endometrium-associated genes segregated samples by hormonal treatment rather than donor variability (N=2 donors), and adequately reproduced gene expression profiles associated with their respective menstrual cycle phases (Fig S9A). To identify the molecular functions, biological processes, and pathways differentially regulated by the hormone treatments, we performed Gene Ontology (GO) analysis and detected key reproductive biological processes associated with progesterone signaling to be significantly (p>0.05) enriched in secretory and menses groups, including ECM composition and receptor signaling activity (Fig 4E). Gene network visualizations of these pathways confirmed that, in response to hormone treatment, the co-cultures reproduced progesterone-induced expression of specific secretory phase associated genes including PRL, PAEP, and ECM-associated genes and downregulation of CPM, MMP9, and cytokine-mediated inflammatory genes (Fig S9B).

Finally, gene set enrichment analysis (GSEA) further confirmed recapitulation of canonical menstruation-associated hallmarks (Fig S10D), including significant upregulation of senescence, inflammatory^80^, and hypoxia^85^ in the withdrawal treatment group compared to E or PE (Fig 4F). To expand on these transcriptomic findings, we also observed expression of matrix metalloproteinases (MMPs), including those primarily produced by either the stromal (e.g., MMP-2) or epithelial populations (e.g., MMP-7, MMP-26).^10, 86^ To corroborate these transcriptomic findings, we also measured temporal changes in matrix metalloproteinases (MMPs) from the spent media (Fig S14C). Together, we co-cultures recapitulated similar temporal expression trends that are observed *in vivo*^87, 88^, specifically a reduced MMP-1, -3 and - 10 expression in the secretory phase compared to proliferative group and reduced MMP-2 compared to the menses group (Fig S10C).

Finally, we experimentally validated the co-culture model using functional assays to assess the morphologic and biochemical changes observed across the human menstrual cycle. First, we examined whether the model captured morphologic features of secretory endometrial glands in response to hormonal stimulation observed *in vivo* (Fig 5A). Progestin treatment caused changes in EEO morphology characterized by thickening of the pseudo-stratified columnar epithelial layer and increased epithelial invaginations (Fig 5B, Fig S10A). Moreover, epithelial secretory function was demonstrated by positive staining of progestogen-associated endometrial protein (PAEP), a marker of the secretory epithelium, in the lumens of the organoids treated with progestin (Fig 5C; Fig S10B). Similarly, a robust induction of stromal decidualization in response to progestin also occurred in the co-cultures by day 15 of the experiment resulting in increased secretion of PRL in the secretory groups and reduced in the menses group (Fig 5D). Moreover, the characteristic negative regulation of PGR during the secretory phase compared to the proliferative phase observed *in vivo*^89, 90^ was demonstrated via immunostaining in both epithelial and stromal populations *in vitro* (Fig 5E). In accordance with literature observations^6^, the temporal analysis across the 15 days showed that co-cultures maintained in baseline E2 conditions showed no significant increase in PRL (Fig 5F), but co-cultures treated with the progestin MPA showed a significant increase in PRL production starting 9 days after MPA treatment initiation (Fig 5F). Finally, we also observed a significant (p=0.031, N=3) doubling of apoptosis (Fig 5G-H; Fig S11) in epithelial glands of the co-cultures in the menses groups (3.06 Casp^+^/cm^2^) compared to the secretory groups (1.37 Casp^+^/cm^2^) mimicking the increased cell death observed during the induction of menstrual phase in vivo.^91, 92^

**Figure 5.**
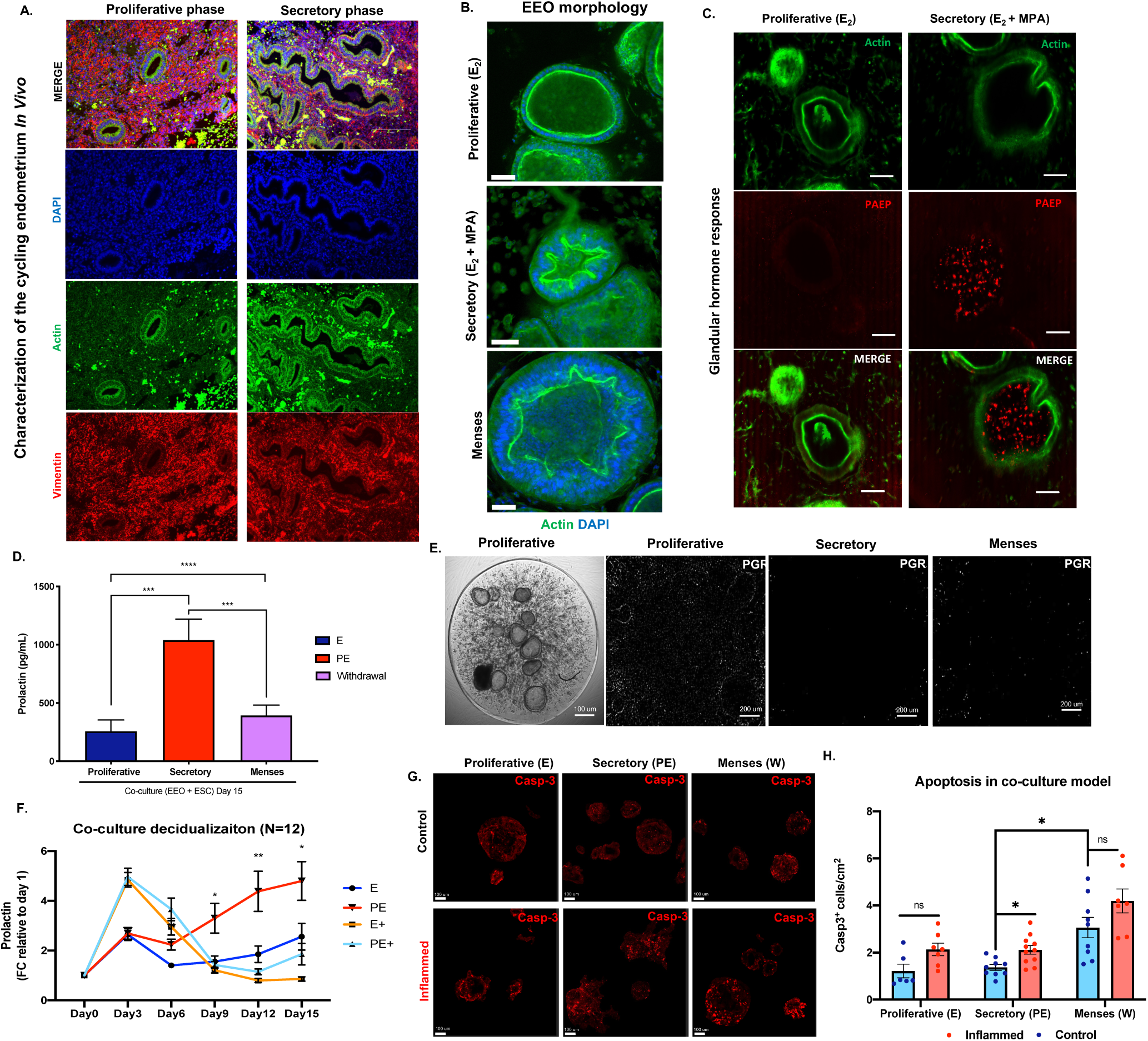
Functional validation of the co-culture model recapitulates morphologic and biochemical cycle-dependent reproductive processes. (A) Representative histological (IF) characterization of the proliferative and secretory phases of the menstrual cycle in vivo reveals morphologic and biochemical changes. Vimentin (red), F-Actin (Green), DAPI (blue). (B) Characterization of the epithelial morphology in co-cultures at day 15 of culture by IF reveals glandular maturation and invagination of EEOs in response to progestin treatment groups. (C) IF staining of PAEP (glycodelin) in co-cultures recapitulates the epithelial secretory phenotype. (D) Stromal decidualization was assessed by measuring prolactin secretion in the medium across 15 days of culture. PRL secretion is induced by PE treatment but suppressed in menses treatment group. (E) PGR staining in the co-cultures at day 15 of treatment. (F) Temporal profiles of decidualization normalized to fold change relative to day 0 were performed by measuring spent media (N=12) and can be suppressed by IL1β treatment (1 ng/mL, E+, PE+). (G-H) Cleaved caspase-3 (Casp-3) IF in the co-cultures as a marker of apoptosis (G) and quantified in as positive staining per organoid area (H). All images are of day-15 co-cultures in the synthetic ECM 3wt% 20kDa-PEG-Mix hydrogels. Significance is indicated as *p<0.05, **p<0.01, ***p<0.001.

Altogether, these results demonstrate the establishment of a co-culture model of the endometrium can recapitulate *in vitro* several of the hormone-induced phenotypic, morphologic, and biochemical changes associated with the idealized human menstrual cycle and could be used to investigate on reproductive function in mechanistic fashion.

#### 3.2.4. Progesterone action is disrupted by IL-1β treatment in the co-culture model

The suppression of stromal cell response to progestin in the presence of IL-1β (Figs 3H,J) motivated us to analyze the molecular and phenotypic responses of epithelia and co-cultured stromal and epithelial cells to IL-1β, anticipating outcomes in the latter case would likely be influenced by stromal-epithelial cross talk. Separate proliferative, secretory, and menses treatment groups were co-stimulated with IL-1β (1 ng/mL) throughout the 15-day experiment (Fig 4D). First, in accord to the results described above (Fig S6), the stromal population in co-cultures also developed an elongated (inflamed) morphology (Fig S15). Furthermore, co-cultures exposed to the inflammatory stimulant IL*-1*β showed suppression of a canonical decidualization response, production of PRL after extended (week or more) exposure to a progestin, as in E (Fig 5F). Whereas the control co-cultures showed a canonical increase in production of PRL following a week of progestin exposure and continued increase through the 15 days of treatment (Fig 5F), PRL production in control co-cultures treated with IL-1β showed no difference in PRL production in the presence or absence of progestin; both E and E+P co-cultures treated with IL-1β showed an unexpected spike in PRL production in the immediate 3 days after stimulation with the inflammatory cue followed by a fall to negligible levels for the remainder of the 15 days (Fig. 5F). Analysis of the inflamed (IL-1β treated) co-cultures by bulk transcriptomic (Fig S13) and multiplex immunoassay (Fig S14) analysis confirmed the decreased PRL expression observed at the protein level (Fig S13B). Gene expression analysis in a generalized linear model using 8,423 genes expressed across all samples comparing controls to IL-1β stimulated cocultures (Fig S13A) revealed an expected global up-regulation of several additional pro-inflammatory cytokines and chemokines (e.g., IL6, TNF-α, CCL2, CCL8, CCL5) and down-regulation of anti-inflammatory cytokines (e.g., IL10) in the IL-1β -treated groups (Fig S13B). Hallmark pathway enrichment analysis of the unstimulated and IL-1β- stimulated co-cultures detected disruptions of progesterone-regulated pathways including the downregulation of ECM structural constituent, and upregulation of inflammatory receptor-ligand signaling processes (Fig S13C). Immunostaining revealed that co-cultures with IL-1β, compared to the untreated controls, further increased apoptosis in the transition from the secretory to the menses phase (Fig 5H; Fig S11), an observation that was corroborated by a separate assay of cell death (Fig S12). These findings show that IL-1β can disrupt progesterone actions. We confirmed this via immunostaining for PGR expression in the co-cultures which showed reduced PGR signal, even in the E-containing media, (Fig 6A) in both the epithelial and stromal populations.

**Figure 6.**
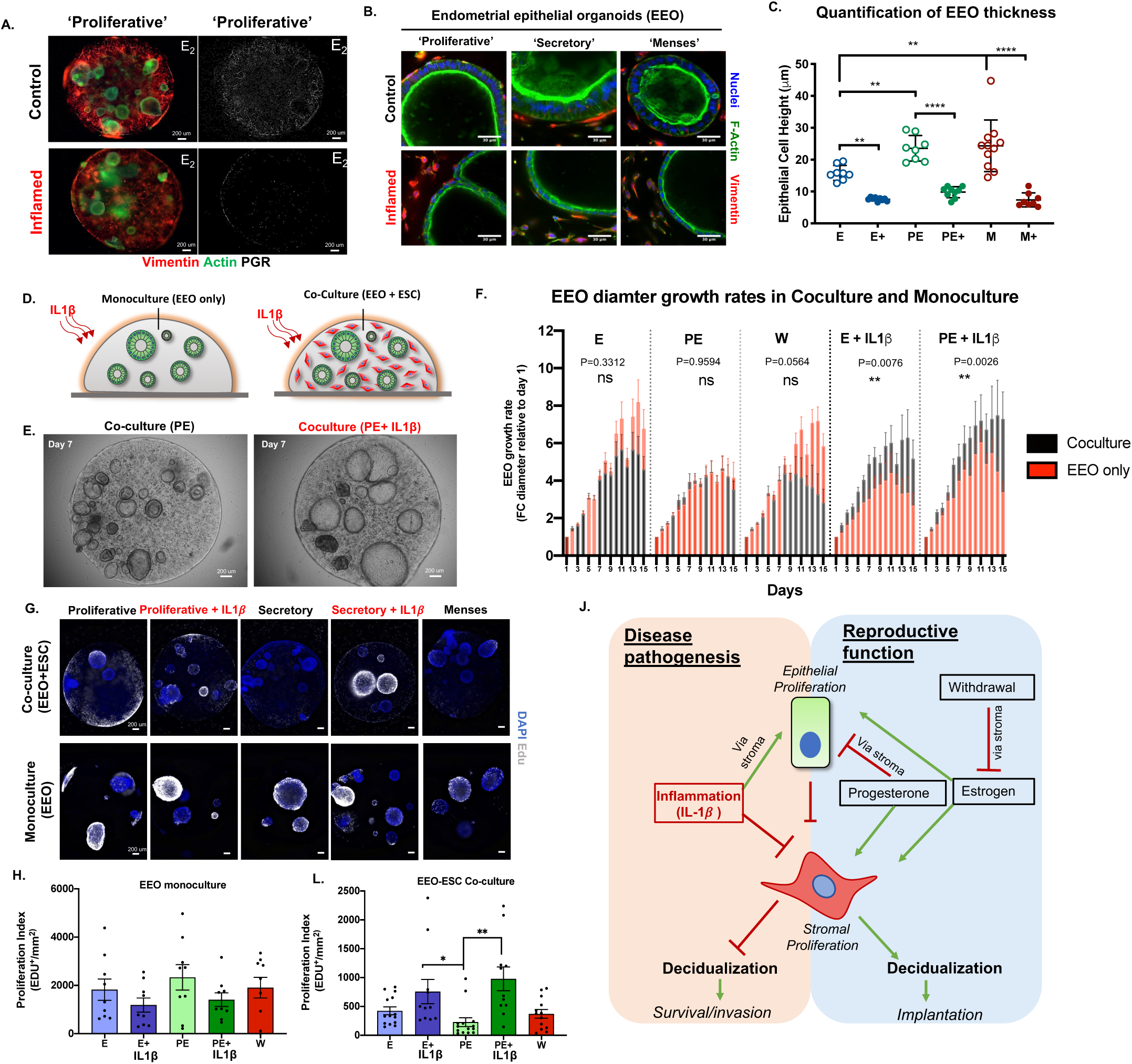
Functional analysis of inflammatory cue (IL-1β) repsonses in co-cultures in synthetic hydrogel reveals the intitiation of the endometriotic phenotype mediated by cell- cell communication. (A) 3D maximum intensity projections of PGR expresison in E2-only treated co-cultures in response to IL-1β stimulation (B-C) IL-1β stimulation (1 ng/mL) induces morphological changes in EEOs. Representative IF images (B) and quantification (C) of epithelial height (distance from base to apical edge of the epithelium, N=3). (D) Schematic of donor matched EEO monoculture and co-culture in IL-1β treatment groups. (E) Represenative images of time-lapse co-cultures stimulated with IL-1β reveals enlarged EEO diameter by day 7 of culture. (F) Quantication of the mean fold-change of EEO diameter across 15-days of cultures from daily time-lapse images in co-cultures (black) and monocultures (red) across treatment conditions (E, PE, W) and IL-1β-treated (E+, PE+) groups. Fold-change denotes the mean increase in diamter relative to day 1 of culture. Analysis compares the mean EEO growth rate between monoculture vs coculture (n=4; E + IL-1 p=0.0076; PE + IL-1β p=0.0026). (G) Immunostaining analysis of DNA synthesis in co-cultures and EEO monocultures by EdU incorporation in response to hormones and IL-1β treatment (N=3). (H-I) Quantification of proliferative profiles calculated as the number of EdU^+^ cells per organoid area (mm^2^) in (H) organoid monocultures and (I) co-cultures. (J) Working model of the cellular mechanisms driving epithelial and stromal communication in the endometrium under physiologic (hormones) and pathogenic (IL-1β -treated) conditions. All images are of day-15 co-cultures in the synthetic ECM 3wt% 20kDa-PEG-MIX hydrogels. Significance is indicated as *p<0.05, **p<0.01, ***p<0.001.

Lastly, stimulating the co-cultures with IL-1β caused a significant flattening of epithelial morphology causing a shift from the pseudo-columnar architecture in the controls toward a squamous cell structure with thin apical F-actin staining (Fig 6B). Whereas progestin treatment significantly increased cell height in the control co-cultures, as characterized by the distance from apical to basolateral side (E vs PE; p=0.0098), in the IL-1β -treated groups, epithelial cells were thinner than in controls (PE vs PE+IL-1β; p <0.0001) and were relatively unaffected by progestin (Fig 6C). This flattening phenomenon closely mimics the de-differentiation events observed in the wound repair cell response of the intestine.^93^ The change in morphology of the epithelial cells was accompanied by a slight, but non-significant (p=0.127), increase in spacing between nuclei compared to the round nuclei in the controls at the single cell level (Fig 6B, Fig S16A), suggesting the flattened cell in the IL-1β treated groups have a slightly increase in projected cell area compared to the controls to compensate for the reduction in cell height while keeping similar volume. An alternate metric of projected cell area, the density of nuclei counted per area of organoid monolayer surface, was comparable in control and IL-1β- treated co-cultures (Fig S16B). Further analysis of the morphometric changes cell volume is a topic for a future investigation. Altogether, these results highlight the extent by which IL-1β can promote the overexpression of additional pro-inflammatory cytokines associated with reproductive diseases^94–96^ and can potentiate progesterone resistance, a phenomenon that is a characteristic feature of endometriotic disease *in vivo*^97, 98^.

### 3.3 IL-1β treatment disrupts epithelial proliferation via a stromal-dominant communication

Chronic inflammation of ectopic lesions is a characteristic feature of endometriosis; wherein endometrial epithelial and stromal cells are found growing outside the uterus.^99^ We thus next assessed whether inflammation altered epithelial proliferation in ways that may illuminate mechanisms of disease. Tissue-recombinant murine models have previously demonstrated that estrogen and progesterone mediate epithelial mitogenic processes indirectly through signals from the hormone-responsive stromal fibroblast,^83, 100, 101^ therefore hypothesized that IL-1β may exert its effect via similar mechanisms. To test this, we measured the effects of IL-1β treatment on epithelial growth, in epithelial monocultures and in co-cultures (Fig 6D; N=4 donors, matched cultures), using both morphological analysis of organoid diameters and measuring DNA synthesis.

First, we measured changes in organoid size (outer diameter) across the 15 days of culture as a proxy for growth rates (Fig S17A). As observed with endometrial stromal cells in monoculture (Fig 3G-H), IL-1β treatment significantly (PE vs PE+IL-1β; p=0.0121) suppressed the growth of the epithelia in monoculture, compared to controls, using the metric of organoid diameter (Fig S17A). Unexpectedly, in co-culture with stroma, IL-1β treatment caused an observable increase in epithelial organoid diameter compared to untreated controls (Fig 6E). Quantification of the organoid diameter across the 15 days of culture demonstrated that IL-1β stimulation positively impacts scEEO diameter growth in the co-cultures, but not in the monocultures (Coculture vs monoculture; p=0.0026) resulting in a mean fold change in organoid diameter of 7.3 and 3.9-fold in the co-cultures and monocultures, respectively (Fig 6F).

As scEEO diameter is only a proxy for changes in cell number, the mitotic activity of these cultures at day 15 was examined by measuring EdU incorporation by the organoids after 24 hours of incubation (Fig 6G). In organoid monocultures there was a non-significant (p > 0.05) trend toward reduction of DNA synthesis in scEEOs upon treatment with IL-1β (E=1822; PE=2333; E+IL-1β=1185; PE+IL-1β=1407 EdU^+^/mm^2^; N=3) and a surprising trend toward increase in DNA synthesis in the presence of progestin compared to E-only conditions (Fig 6H). In stark contrast, stimulation with IL-1β resulted in a greater than 3-fold increase (PE = 228; PE+IL-1β = 978 EdU^+^/mm^2^; P= 0.0014; N=4) of scEEO DNA synthesis in cocultures compared to controls (Fig 6I), consistent with the observed increase in organoid diameters in IL-1β treated co-cultures (Fig 6E). Further, unlike monocultures, the proliferative index of scEEOs in co-culture decreased in progestin-treated controls compared to E-only controls, as would be expected in the transition from proliferative to secretory phase *in vivo*^102^. Unexpectedly, the proliferative index of scEEOs in co-cultures treated with IL-1β showed the opposite trend to that of monocultures when progestin was added– an increase in the proliferative index compared to E-only cultures (Fig 6E). These phenotypic changes in epithelial mitotic activity mirror the increased epithelial proliferative profiles observed in the eutopic and ectopic endometriotic tissues in vivo.^103, 104^ Although the scEEO monocultures appeared to have a greater proliferative index than the co-cultures in some groups, this difference could be attributed to the contribution of the ESC in mediating hormone signaling, as shown in other models.^105^

Perhaps because of this increased growth rate, IL-1β stimulation in co-cultures resulted in greater organoid collapse by day 15 and could also explain the ∼1.5-fold (p=0.012) increase in apoptosis (Casp-3^+^ staining) observed in the epithelial (Fig 5G) populations that resulted in an overall increase in cell death (ethidium homodimer staining, Fig S12). Altogether, these results support a working model by which inflammatory signals and sex hormone signaling in the endometrium are partially mediated indirectly via the stroma (Fig 6J).

## Discussion

The endometrium is a marvelous example of regenerative biology wherein sex hormones mediate rapid growth and maturation of the tissue, accompanied by equally dynamic changes in ECM-cell interactions that are both mechanically and molecularly linked to reproductive function.^106^ These remarkable regenerative properties also contribute to common debilitating diseases like endometriosis, for which new therapies are desperately needed. Here, we developed a new model for dissecting molecular and phenotypic consequences of endometrial epithelial-stromal crosstalk in long term (15 day) cultures of patient-derived endometrial cells by defining a completely synthetic extracellular matrix hydrogel that is tailored to both replace Matrigel for organoid culture, and simultaneously to support stromal culture. We then used this model to show how the hormone-dependent behaviors of the endometrial epithelium in co-culture with stroma diverge from those in monoculture, observing for example that the pro-inflammatory cue IL-1β appears to drive the endometrial co-cultures, but not monocultures, to a state that phenocopies features of diseases like endometriosis.^95, 99^

The synthetic ECM described here overcomes certain limitations of organoid technologies and other co-culture approaches for parsing cell-cell communication in general, especially in the endometrium. Natural matrices like Matrigel and collagen – which have been used either alone for co-culture^27, 28, 42, 107^ or combined in creative ways^108, 109^ – include many extraneous signaling molecules that may drown out the signals produced by the cells they support. This limitation is underscored by the inability to design experiments that require longer term stable cultures, as is the case for hormone signaling, or need specific ECM stiffness. The main constituent of this synthetic ECM, PEG, is a blank slate known for its relative lack of interaction with proteins.

Thus, growth factors, cytokines, and other molecules produced by each cell type can freely dominate the cell-cell communication networks. By design, the synthetic ECM comprises only a minimal set of biological cues: two integrin ligands, two ECM-binding proteins, and a peptide crosslinker, all produced by standard, well-defined chemical peptide synthesis methods. These minimal cues, which were modified from a previous formulation used for organoids^26^ and monolayer cell line co-cultures^25^, include ligands for a spectrum of integrins expressed differentially by epithelial and stroma (see Results) and enable cell-mediated remodeling of the microenvironment as well as the accumulation of cell-produced ECM. By demonstrating that this synthetic matrix is sufficient to generate scEEOs from the original primary glands without the need to first expand in Matrigel (Fig S18), we provide a hydrogel alternative to alleviate the dependency on Matrigel. Because the synthetic matrix is relatively simple and is chemically defined, it is straightforward to implement reproducibly for the culture of organoids and co-cultures. Reproducibility within and between labs is highly desirable to analyze phenotypes of patient-derived tissue models due to the inherent biological variability in the samples themselves. Finally, although only an initial characterization is provided here, the synthetic ECM arguably offers greater potential to mimic and study certain features of endometrial diseases compared to ECM-free co-culture models^110, 111^, as its biophysical properties can be tuned systematically to mimic the “soft” regimes of healthy endometrium or the “stiff” regimes of the myometrium^112^ or fibrotic tissue (Fig 2A).

Steroid hormone-driven stromal-epithelial crosstalk in the endometrium has been illustrated in mouse models involving cell type-specific receptor knockouts in the eutopic endometria^89, 113^ and recombinant implantation of tissue fragments in the renal capsule,^105, 114^ but crosstalk in normal and diseased human co-cultures has been more challenging to parse due to the disparate matrix environments preferred by each cell type^115, 116^, and the relative short-lived nature of cultures in Matrigel or collagen. For example, work by Rawlings and co-workers showed stromal-epithelial crosstalk enabled a simplified medium to support an elegant collagen gel-embedded “assembloid” co-culture model that captured both normal and drug-skewed co-evolution of the stromal population to subpopulations of decidual and acutely senescent fibroblasts^42^. They thus defined a tantalizing model for analysis of embryo implantation if the limitations of the culture longevity, which disintegrate before the entire sequence of implantation steps is realized under some conditions, and the limited access organoid apical interface, can be overcome^42^. Thus, the long-term capacity of co-cultures in our synthetic hydrogel may provide one approach to extending the assembloid model if similar evolution of fibroblasts populations can be demonstrated. In concordance with a different short term (7-day) co-culture model, employing EEOs cultured in Matrigel in a Transwell membrane above a coverslip coated with stroma^117^, we observed the expected suppression of proliferation by E2 and P4 compared to E2 in control co-cultures (Fig 6F and Fig S17). However, we are also able to observe more pronounced effects at the two-week time point, especially in cultures treated with IL-1β (Fig 6F and Fig S17) and provide evidence that the inflammatory cue IL-1β can also modulate epithelial proliferation via the stroma (Fig 6J).

Indeed, the responses of co-cultures to inflammatory cues at extended time points are the most striking findings in this study, as they may shed light on endometrial disorders closely associated with pro-inflammatory cytokines such as TNF-α and IL-1β.^94^ In co-cultures treated with IL-1β, organoid diameter (Fig 6F) and epithelia mitotic activity (Fig 6I) are both increased, even in the presence of MPA, compared to the E2 condition, whereas the canonical response to progestin is a decrease in epithelial proliferation, as observed in the control co-cultures and in monocultures. We speculate that this observation of an IL-1β-MPA synergy may have clinical relevance for the observed failure of some endometriosis patients to respond to progestin therapies. Similar to our observations about IL-1β inducing profound epithelial flattening along with increased proliferation, in deep infiltrating endometriosis lesions, an “invasive front” characterized by highly flattened epithelia with greater proliferative index than epithelia in a more quiescent core has been described^118^. Furthermore, greater proliferation indices are observed in the eutopic endometrium of endometriotic patients, compared to controls^102, 104^. While more translational validation is warranted to confirm mechanistically similar behaviors, the results are suggestive that the model may capture features that are important clinically.

Importantly, the timely and acute induction of inflammatory mediators are also well known to be critical for maintaining normal reproductive physiology^79^. This may explain why a transient (day 3) increase in PRL secretion was observed in the co-cultures in response to IL-1β as acute-phase inflammatory insults that activate the nuclear factor–kappaB (NF-κB) system, a master regulator of cell survival and proliferation in endometriotic disease^119^, have been shown to regulate PGR expression in vivo^79, 89^, and can enhance stromal decidualization and blastocyst implantation^55, 120^ A limitation of this study is that most assays were performed at end point measurements (day 15), future mechanistic studies that dissect the timing (acute vs chronic), abundance (low vs high concentration), and synergism with sex hormones (estrogenic vs progestin conditions) will be critical to fully understand the extent by which IL-1β can positively promote physiologic (e.g., implantation) or induce pathologic processes (e.g., endometriosis, infertility)^98, 119^, and ultimately will help identify therapeutic approaches for targeting the disease while maintaining reproductive function (Fig 6J).

Finally, we are just beginning to explore how the properties of the synthetic ECM may enable more complex tissue-level architectures than the relatively simple organoid-stroma culture described herein. For example, we have observed that organoids close to the gel-liquid interface sometimes merge with each other and erupt to form mature epithelial organoid- monolayers structures that are more representative the glandular-luminal epithelial compartment of a true endometrial mucosal barrier (Fig S21). We are now defining protocols that foster efficient and reproducible fusion. A parallel effort to represent the heterogenous cell populations of the endometrial microenvironment combines immune cells with stroma and epithelia, as we have described for pancreatic tumors^31^, with an eye toward addition of these complex endometrial models to perfusable blood vessels maintained in microfluidic devices, extending our initial findings that we can form microvascular networks in synthetic PEG gels^121, 122^. We speculate that the co-culture model described here can be productively extended by the research community as a platform to mechanistically parse stromal-epithelial crosstalk in the human endometrium.

## Acknowledgements

We thank the Newton Wellesley Hospital staff for facilitating a tissue collection program with MIT and the patients that donated their tissues for our research. We thank Histology Facility at the Koch Institute at MIT for their assistance with histological processing of tissues. We thank Profs. Kevin Osteen and Kaylon Bruner-Tran at Vanderbilt University for their support. J.S.G. was supported by an Environmental Toxicology Training Grant (NIH T32 ES007020) and from the Bill and Melinda Gates Foundation. Additional funding was provided by The John and Karine Begg Foundation, the Manton Foundation, and NIH U01 (EB029132).

## Author Contributions

J.S.G. and A.T.B. conceived the study, designed, performed, and analyzed the experiments. J.S.G. wrote the manuscript. J.S.G isolated, cultured and maintained primary cell populations with the help of C.I. and L.B. In Vitro culture data processing and analysis was performed by J.S.G., A.T.B., and K.B. Hydrogel and tissue biobank was maintained by V.H.G, and C.I. Ex Vivo analysis was performed by B.G. and L.B. for scRNAseq and proteomics, respectively. All were involved in editing the manuscript. M.L., K.I., and J.S.G. supervised tissue collection and processing. L.G.G. supervised project development and edited the manuscript.

## Data availability

Source data are available upon request from the corresponding author.

## 6. Supplementary Figures Legends

**Supplementary Fig 1 . Characterization of the human endometrium using single cell transcriptomic analysis of the endometrium.** (A) Expression of all integrin chains across the cycle for diverse subpopulations of endometrial cells (N=6); integrin β3 was not detected. (B) Violin plots of the integrin expression across the endometrial cell populations (N=6). (C) Validation of cell specific integrin expression in cultured endometrial cells. ESC and EEOs were cultured in 2D polystyrene (ESC) and in Matrigel (EEO) respectively and analyzed by qPCR for integrins relative to GAPDH (N=5). Significance is indicated as *p<0.05, **p<0.01, ***p<0.001.

**Supplementary Fig 2. Workflow for EEO generation, hydrogel generation and screening.**

(A) Schematic of primary endometrial tissue processing workflow to generate organoids (B) Schematic of detailed EEO generation and hydrogel screening protocols.

**Supplementary Fig 3. Screening and quantification of EEO generation in synthetic matrices across donors.** Representative brightfield images of scEEO expansion comparing 3 different donors at day 7 cultured in either Matrigel or one of 4 different hydrogel formulations.

**Supplementary Fig 4. Matrix stiffness affects EEO morphology.** (A) Represenative brightfield images of day-14 scEEO morphology donors cultured in hydrogel formulations of GFOGER (3mM) or MIX (1.5 mM GFOGER, 1.5 mM PHSRN-K-RGD) in varying stiffnesses (3wt%, 5wt% and 7wt%). (B) Quantification of meandering phenotype as a percentage of EEOs confering a meandering phenotype (N = 4 biological replicates). (C) Representative maximum intensity projection and orthogonal view of scEEOs in Matrigel, soft PEG MIX gel, and stiff PEG MIX gel, stained for F-Actin (green).

**Supplementary Figure 5. Comparative phenotype, proliferative activity and heterogenity of EEOs cultured in PEG vs Matrigel.** (A) Representative image of F-actin stained scEEO derived in soft synthetic matrices compared to Matrigel at day 14 reveal a comparable rounded spherical morpholgy. (B) EpCAM staining of scEEOs confirms epithelial-origin in synthetic matrices and in Matrigel. (C) Ki67 staining of EEOs cultured in synthetic ECM and in Matrigel. (D) Image captures from videos reflect epithelial heterogeneity and motile cilia. EEOs cultured in synthetic matrices and stained for acetylated a-tubulin (red) reveal multi-ciliated and primary ciliated cells.

**Supplementary Fig 6. Inflammation disrupts ESC morphology.** (A) Representative brightfield images of ESC monocultures in response to inflammatory cue (IL-1β, 1 ng/mL), reveals increased elongation suggestive of a more active and motile phenotype. (B-C) Imaging and quantification of increased ESC remodeling capacity. (B) Immunofluorescent characterization of F-actin confirms morphogenic changes associated with response to IL-1β treatment (4X). Single cell segmentation (False colored red, 20X) cells elucidates the elongated morphologic changes associated with hormone treatment and IL-1β treatment in 3D (N=3). (C) Quantifaction of mean stromal cell surface area (N=3).

**Supplementary Fig 7. Estimating cell ratios ex vivo for tissue modeling in vitro.** 3D imaging of endometrial tissue biopsies using tissue clearing methods and confocal imaging were analyzed using the Imaris (Bitmap 9.6.0) software to segment the epithelial and stromal compartments of the native endometrium (n=3 donors; prolifertive n=2, secretory n=1; 3 different volumes) using the ‘Surface’ function to masks individual cell regions in a volumetric image. (A) The ‘Cells’ function Single cell segmentation tools from the software were used to calculate the number of epithelial cells (Epi) per volume of epithelial glands (Epi/µL). (B) ESC single cell counting in 3D was performed using the ‘Spots’ function by labeling and counting the DAPI+ nuclei in the image. This estimations is calculated as the stroma, including other cellular components including the vascular and immune components. A similar approach was taken for co-culture in the *in vitro* models. (C) Cell concentration per unit area the native endometrium and the cell ration between epithelial cells and stromal cells. (D) 3D analysis of EEOs cultured in the synthetic matrices. (E) Cell density in vivo and in vitro comparing relative numbers and epithelial cell gland volumes. (F) No significant differences in the ratio between single ESCs concentration (ESC/µl) and Epithelial volume (gland/volume ratio).

**Supplementary Fig 8. Establishment and transcriptomic analysis of monocultures and co- cocultures.** (A) Representative brightfield images of co-culture (N=7) and monocultures of EEOs (N=1) and ESCs (N=6). (B) Unsupervised PCA analysis of bulk RNAseq show that day-15 co-cultures and monocultures cluster by culture type. Ellipses bound the 95% confidence interval for each group. (C) Plotting *log*_2_(*fold changes*) against −*log*_10_(*P value*) in a volcano plot shows differentially expressed genes in cocultures as compared to cultures of ESC only.

**Supplementary Fig 9. Trancriptomic evaluation of endometrial co-culture models of the menstrual cycle phases.** (A) Hierarchical clustering heatmap of genes associated with the human menstrual cycle clusters samples based on hormone treatment groups in the co-cultures. Data is derived from 2 biological replicates (donors) that were maintained in nEEO media (B) Examination gene network analysis of the most significant up-and-down regulated GO associated genes.

**Supplementary Fig 10. Functional and trancriptomic evaluation of endometrial co-culture recapitulation of the menstrual cycle phases.** (A) Morphological assessment of altered cellular response to hormones and to inflammation (IL1β, 1ng/mL). Additional representative images of F-actin staining in EEO (green) and ESC (faux colored, orange) in co-culture under ‘proliferative (E)’ and ‘secretory (PE)’ hormone treatment groups for both EEOs and ESC. (B) Low- magnfication (4X) immunofluorescent staining of PAEP of the day-15 co-cultures reveals progesterone-induced secretory phenotype in the EEOs. (C) Transcriptomic analysis of MMP expression reveals differential expression patterns in response to hormone treatments. (D-E) Gene Set Enrichment Analysis (GSEA) on a ranked list of genes from a generalized linear model of the coculture bulk RNAseq data comparing different treatment groups. Hallmark pathways show gene expression changes consistent with corresponding phenotypic changes across the phases of the menstrual cycle.

**Supplementary Fig 11. Apoptosis is increased in the menstrual phase and with IL-1β treament in the co-culture model** (A) Comparison of apoptotic marker expression (cleaved caspase-3) immunofluorescent staining in co-cultures at day 15 in the menstrual cycle and inflamed groups in low (4X) magnification. E = estradiol only, PE = E + MPA, M = PE withdrawal + RU486), E+ = E + IL1β, and PE+ = PE+IL1β, M+ = M + IL-1β.

**Supplementary Fig 12. Cell viability is decreased in the IL-1β treatment group in the co-culture models at day 15 in the 3% PEG MIX hydrogel.** Low (4X, scale bar = 100 ums) and high (10X, scale bar = 50 ums) magnification of the Live (green) and Dead (red) assay in the hormone and IL-1β-treated groups comparing (A) ESC monocultures and (B) co-cultures in 3 ul droplet cultures.

**Supplementary Fig 13. Co-cultures treated with IL-1β have transcriptomic profiles consistent with inflamed endometrial states.** Co-cultures treated with IL-1β have transcriptomic profiles consistent with inflamed endometrial states. (A) A volcano plot of *log*_2_(*fold changes*) against –*log*_10_(*P value*) for cocultures treated with the inflammatory cue (IL- 1β, 1 ng/mL) as compared to non-inflamed controls. Transcript data is modeled using a generalized linear model (see Methods) to account for inter-donor and hormone treatment variability. (B) A closer examination of the most significant GO terms reveals that canonical inflammatory genes are upregulated while PRL is downregulated, as expected for an inflamed environment. (C) Differentially expressed genes were analyzed using gene ontology (GO) analysis to examine the most up- and down- regulated biological processes upon IL1β treatment.

**Supplementary Fig 14. Temporal evaluation of inflammatory response of co-cultures via targeted proteomics.** Represenative co-culture (N=1 donor) of measured multiplex 45-panel of secreted cytokines measured in co-cultures throughout the 15-day experiment. Readings were measured every 3 days in control or IL-1β-treated conditions (N=3 technical replicates per day). (A) Luminex panel of up regulated and downregulated cytokines in response to IL-1β-treatment (representative results shown for day 15; similar data obtained for all conditions and time points). (B) Principal component analysis (PCA) and corresponding loading plots illustrate separation of culture conditions. (C) Profiles of MMP release into the medium in co-cultures. All images are of day-15 co-cultures in the synthetic ECM (3wt%-MIX). (E = estradiol only, PE = E + MPA, M = PE withdrawal + RU486), E+ = E + IL1β, and PE+ = PE+IL1β, M+ = M + IL-1β).

**Supplementary Fig 15. Inflammatory stimuli alter stromal morphology in endometrial co-culture.** (A) Representative brightfield image of ESC laden region in the co-culture in the PE treated groups stimulated with and without IL-1β at day 10 of culture in EEO co-culture media (B) IL-1β treated stromal cells revealed increased spindle morphology in co-cultures as stained for vimentin (red).

**Supplementary Fig 16. Cell nuclei spacing and cell density in the organoid surface is comparable across treatment group conditions.** A) Quantification of spacing between nuclei in optical sections of the organoids (left) and representative images of treatment groups (right), (N=3 donors). (B) Representative 3D image projection of nuclei distribution of organoids in the treatment groups. Nuclei density quantificaiton of per organoid surface area is comparable between treatment conditions. Image analysis was performed with Imaris software (N=3).

**Supplementary Fig 17. Quantitative assesment of individual EEO diamater growth rates in in co-cultures and monocultures across 15-days of culture.** Daily images were analyzed to quantify the diameter fold change of the organoids relative to day 1 of cultures. (A) Mean EEO growth rates of co-cultures and EEO only conditions across 15 days of culture (B) Analysis of individual day differences between donor matched co-culture (N=4) and monoculture (N=4) for all treatment groups. Data are mean ± SEM of 4 matched replicates. Signficance is indicated as *p<0.05, **p<0.01, ***p<0.001.

**Supplementary Fig 18. EEOs can be generated and expanded directly into synthetic ECM from primary tissues.** (A) Schematic of epithelial gland collection and EEO generation and expansion using synthetic matrix with Matrigel controls in parallel as a benchmark. (B) Brightfield images of EEOs generated in the synthetic matrix (3wt%-MIX) directly from freshly isolated gland fragments. (C) Characterization of Matrigel-free EEOs and immunofluorescent images of actin (green) staining. (D) Illustration of expansion and passaging of EEOs (p1) using the synthetic matrix.

**Supplementary Fig 19. Trancriptomic evaluation of culture media effect on endometrial co-culture.** (A) Hierarchical clustering heatmap of genes associated with the human menstrual cycle clusters samples based on hormone treatment groups in the co-cultures in 5 biological replicates (donors) that were maintained in ‘EEO’ media.

**Supplementary Fig 20. Histological characterization of endometrial tissues collected.**

Hematoxylin and eosin stain (H&E) of endometrial tissue from endometrial donors.

**Supplementary Fig 21. Synthetic matrix enables modeling of endometrial morphogenesis.**

(A) Schematic of the endometrial architecture capturing the glandular (GE) and luminal (LE) epithelium. Experimental workflow of organoid morphogenic events that lead to the generation of glandular-monolayer structures in the synthetic matrix (3wt% PEG-MIX). (B) Schematic and representative images of the sections of the endometrial luminal and glandular epithelium. Immunofluorescent images of the monolayer section (Luminal epithelium, top) and the organoid section (gladular epitheliumm bottom). Cells are stained for Laminin (white), Dapi (blue), F-Actin (green). (C) Orthogonal view projections of z-stack images of the monolayers generated in traditional 2D polystyrene plates (left) and in the synthetic gels (right) reveals a columnar epithelial morphology in the gel monolayer compared to the 2D where the cells have lost their architecture. Cells are stained for F-Actin (green) and DAPI (blue).

**Table 1. Metadata of endometrial tissue donors.** Table outlining patient metadata and tissue processing.

**Video 1A: Beating multi-cilliated cells are visible in endometrial organoids cultured in Matrigel.** Live recording of beating cilia in organoids cultured in Matrigel at day 6 of culture estrogen (E) treatment groups. Recoding was acquired with a Keyence BZ-X800 Fluorescence Microscope recording option at 40X magnification.

**Video 1B: Beating multi-cilliated cells are visible in endometrial organoids cultured in synthetic hydrogels.** Live recording of beating cilia in organoids cultured in synthetic matrix (3wt%-*MIX*) at day 6 of culture estrogen (E) treatment groups. Recoding was acquired with a Keyence BZ-X800 Fluorescence Microscope recording option at 40X magnification.

**Video 2: Immuno-stained 3D rendering of proliferative phase treated organoid co-culture.** Representative 3D visualization of a z-stack (∼635 um thickness) section of the stained endometrial co-culture in the synthetic ECM (3wt%-*MIX*) at day-15 of culture in the ‘proliferative’ treatment group. Images were captured using a ZEISS confocal Laser Scanning Microscope (LSM 880). 3D renderings were performed using Imaris (Bitmap 9.6.0) software to segement the organoids. Epithelium is stained for F-actin (green) and stroma stained for vimentin (red). DAPI was used as a counter stain for the nuclei.

**Video 3A: Time-lapse imaging of the endometrial co-culture in the ‘proliferative’ treatment group.** Representative (donor 259) brightfield time-lapse of endometrial co-culture in the synthetic hydrogel across 15 days of culture in the ‘proliferative’ (E) treatment group. Images were taken daily at 4X magnification with a Keyence BZ-X800 Fluorescence Microscope.

**Video 3B: Time-lapse imaging of the endometrial co-culture in the ‘proliferative + IL1β’ treatment group.** Representative (donor 259) brightfield time-lapse of endometrial co-culture in the synthetic hydrogel across 15 days of culture in the ‘proliferative’ + IL1β (E+) treatment group. Images were taken daily at 4X magnification with a Keyence BZ-X800 Fluorescence Microscope.

**Video 3C: Time-lapse imaging of the endometrial co-culture in the ‘secretory’ treatment group.** Representative (donor 259) brightfield time-lapse of endometrial co-culture in the synthetic hydrogel across 15 days of culture in the ‘secretory’ (PE) treatment group. Images were taken daily at 4X magnification with a Keyence BZ-X800 Fluorescence Microscope.

**Video 3D: Time-lapse imaging of the endometrial co-culture in the ‘secretory+ IL1β’ treatment group.** Representative (donor 259) brightfield time-lapse of endometrial co-culture in the synthetic hydrogel across 15 days of culture in the ‘secretory’ + IL1β (PE+) treatment group. Images were taken daily at 4X magnification with a Keyence BZ-X800 Fluorescence Microscope.

**Video 3E: Time-lapse imaging of the endometrial co-culture in the ‘menses’ treatment group.** Representative (donor 259) brightfield time-lapse of endometrial co-culture in the synthetic hydrogel across 15 days of culture in the ‘menses’ treatment group. Images were taken daily at 4X magnification with a Keyence BZ-X800 Fluorescence Microscope.

**Video 3F: Time-lapse imaging of the stromal monoculture in the ‘proliferative’ treatment group.** Representative (donor 259) brightfield time-lapse of endometrial ESC monoculture in the synthetic hydrogel across 15 days of culture in the ‘proliferative’ (E) treatment group. Images were taken daily at 4X magnification with a Keyence BZ-X800 Fluorescence Microscope.

**Video 3G: Time-lapse imaging of the stromal monoculture in the ‘proliferative + IL1β’ treatment group.** Representative (donor 259) brightfield time-lapse of endometrial ESC monoculture in the synthetic hydrogel across 15 days of culture in the ‘proliferative’ + IL1β (E+) treatment group. Images were taken daily at 4X magnification with a Keyence BZ-X800 Fluorescence Microscope.

**Video 3H: Time-lapse imaging of the stromal monoculture in the ‘secretory’ treatment group.** Representative (donor 259) brightfield time-lapse of endometrial ESC monoculture in the synthetic hydrogel across 15 days of culture in the ‘secretory’ (PE) treatment group. Images were taken daily at 4X magnification with a Keyence BZ-X800 Fluorescence Microscope.

**Video 3I: Time-lapse imaging of the stromal monoculture in the ‘secretory+ IL1β’ treatment group.** Representative (donor 259) brightfield time-lapse of endometrial ESC monoculture in the synthetic hydrogel across 15 days of culture in the ‘secretory’ + IL1β (PE+) treatment group. Images were taken daily at 4X magnification with a Keyence BZ-X800 Fluorescence Microscope.

**Video 3J: Time-lapse imaging of the stromal monoculture in the ‘menses’ treatment group.** Representative (donor 259) brightfield time-lapse of endometrial ESC monoculture in the synthetic hydrogel across 15 days of culture in the ‘menses’ treatment group. Images were taken daily at 4X magnification with a Keyence BZ-X800 Fluorescence Microscope.

**Video 4A: Time-lapse imaging of the EEO monoculture in the ‘secretory+ IL1β’ treatment group.** Representative (donor 272) brightfield time-lapse of endometrial EEO monoculture in the synthetic hydrogel across 15 days of culture in the ‘secretory’ + IL1β (PE+) treatment group. Images were taken daily at 4X magnification with a Keyence BZ-X800 Fluorescence Microscope.

**Video 4B: Time-lapse imaging of the ESC monoculture in the ‘secretory+ IL1β’ treatment group.** Representative (donor 272) brightfield time-lapse of endometrial ESC monoculture in the synthetic hydrogel across 15 days of culture in the ‘secretory’ + IL1β (PE+) treatment group. Images were taken daily at 4X magnification with a Keyence BZ-X800 Fluorescence Microscope.

**Video 4C: Time-lapse imaging of the endometrial co-culture in the ‘secretory+ IL1β’ treatment group.** Representative (donor 272) brightfield time-lapse of endometrial co-culture in the synthetic hydrogel across 15 days of culture in the ‘secretory’ + IL1β (PE+) treatment group. Images were taken daily at 4X magnification with a Keyence BZ-X800 Fluorescence Microscope.

