## Supplementary data for "Organoid co-culture model of the cycling human endometrium in a fully-defined synthetic extracellular matrix reveals epithelial-stromal crosstalk"

### Supplementary Figures

**Supplementary Fig 1. Characterization of the human endometrium using single cell transcriptomic analysis of the endometrium.**

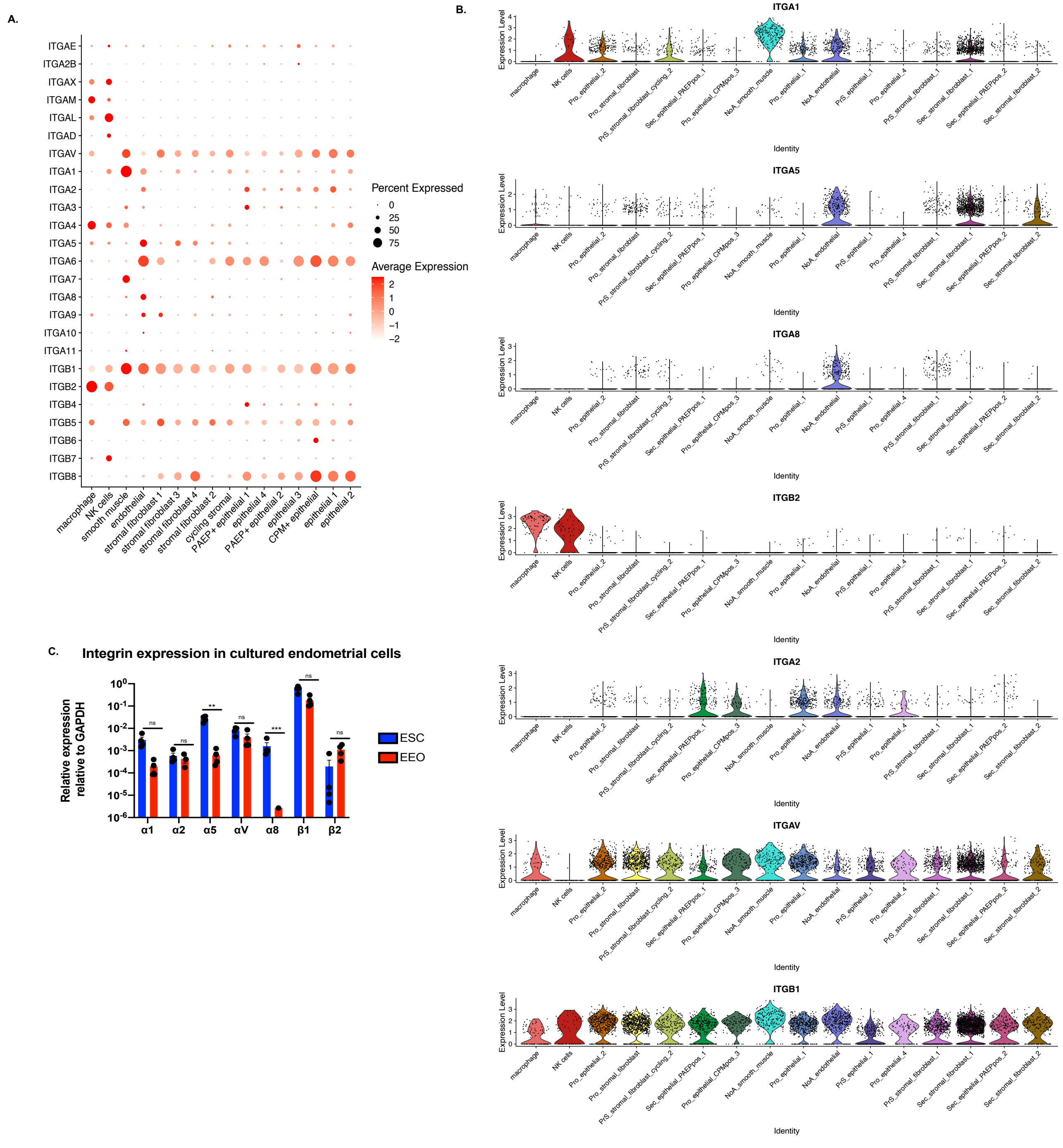

**Supplementary Fig 2. Workflow for EEO generation, hydrogel generation and screening.**

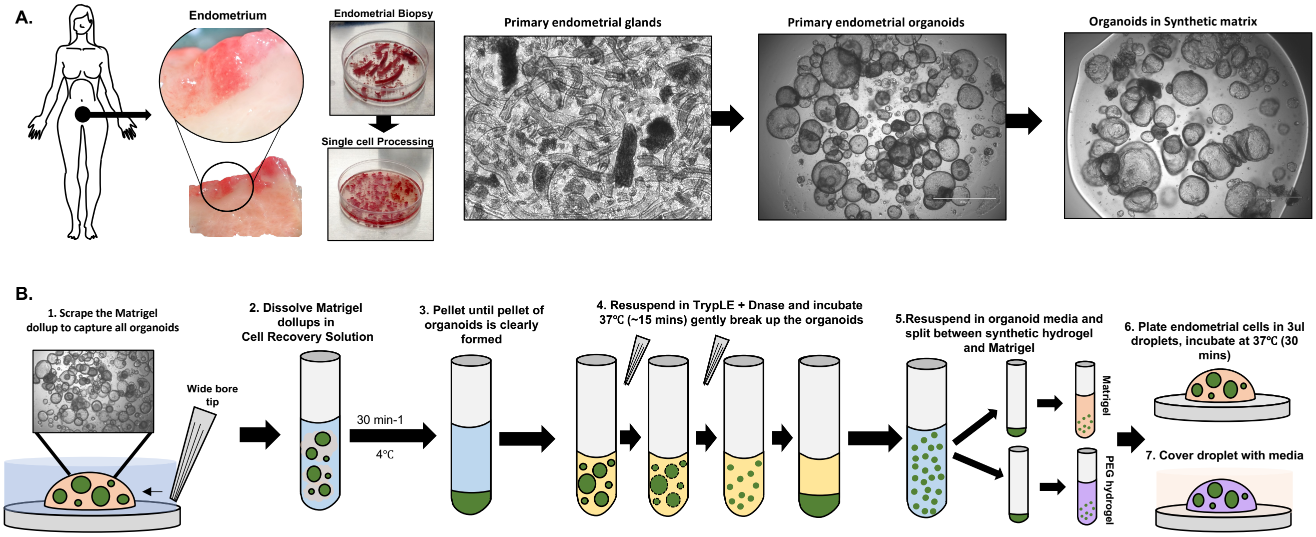

**Supplementary Fig 3. Screening and quantification of EEO generation in synthetic matrices across donors.**

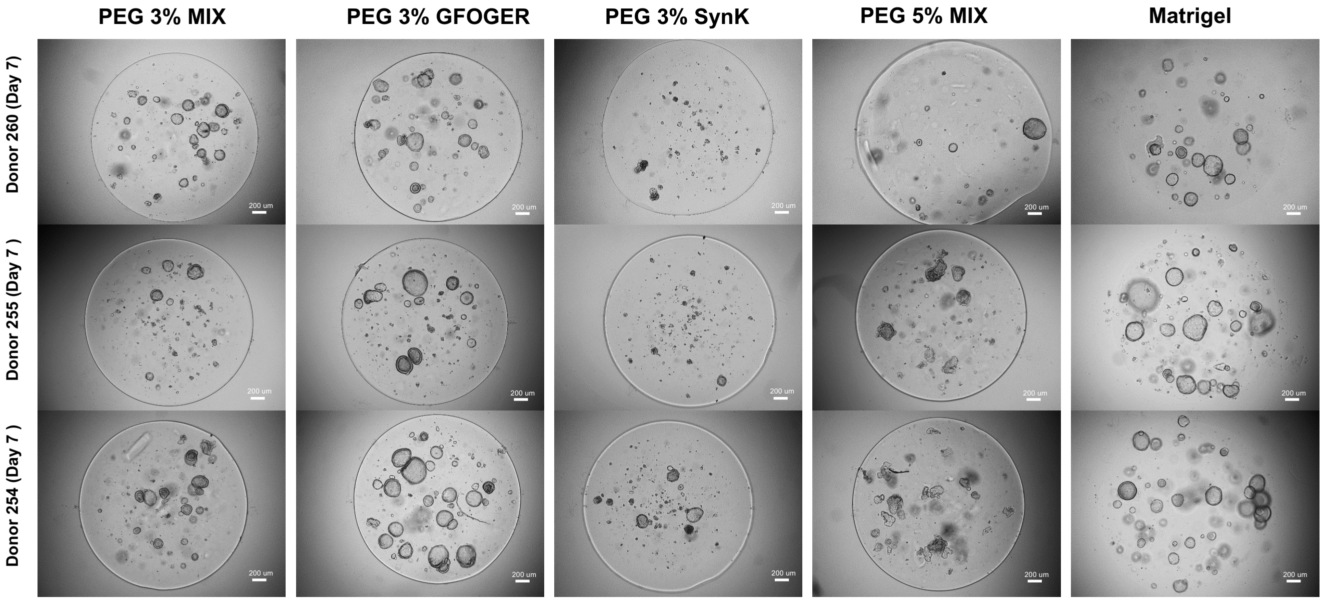

### Supplementary Fig 4. Matrix stiffness affects EEO morphology

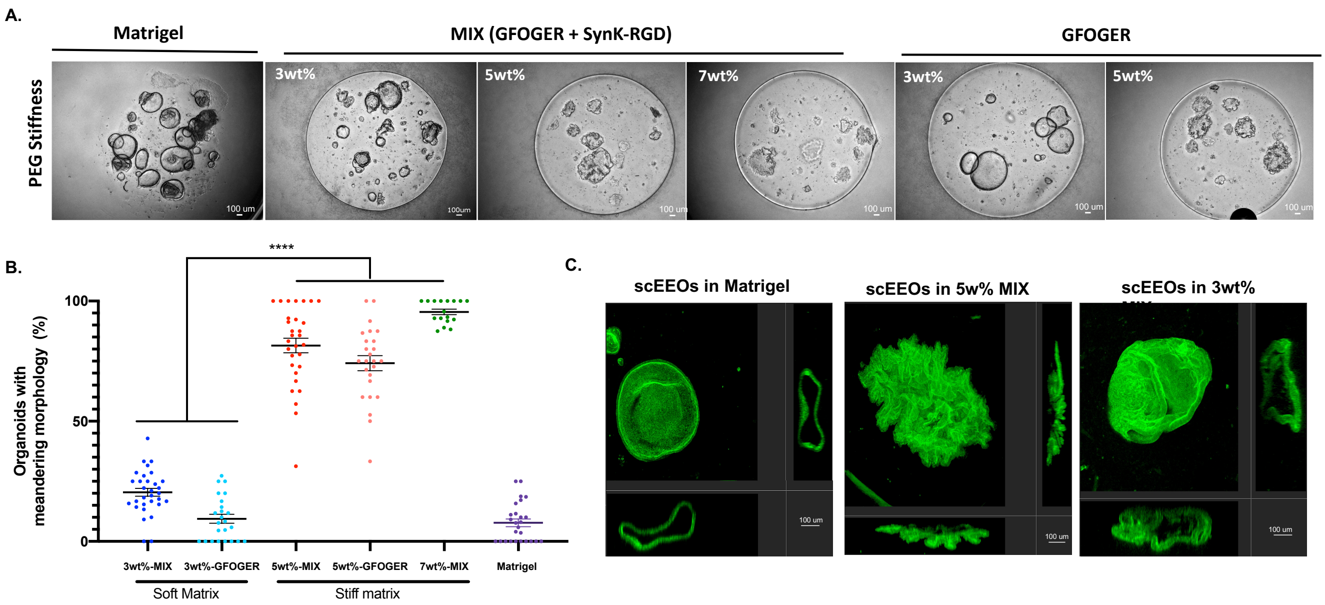

**Supplementary Figure 5. Comparative phenotype, proliferative activity and heterogenity of EEOs cultured in PEG vs Matrigel.**

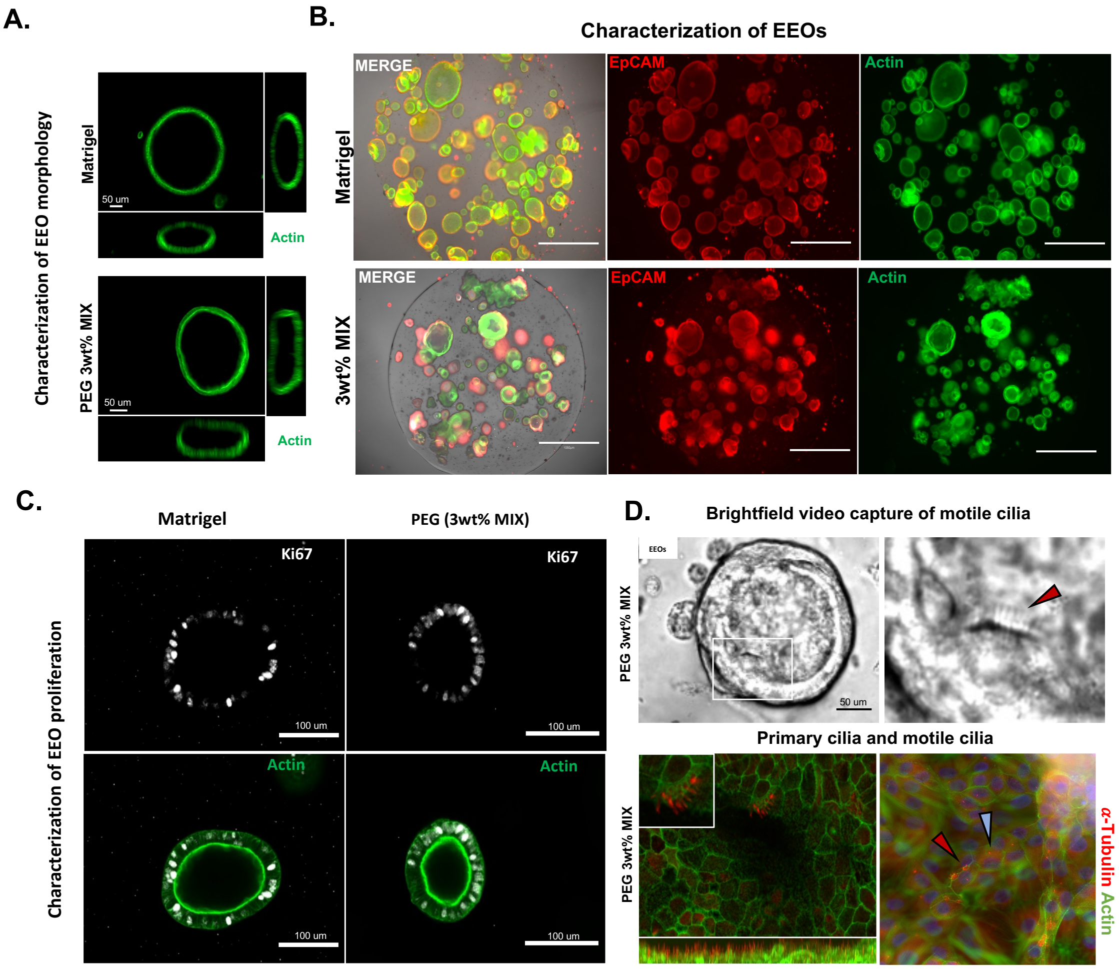

#### Supplementary Fig 6. Inflammation disrupts ESC morphology.

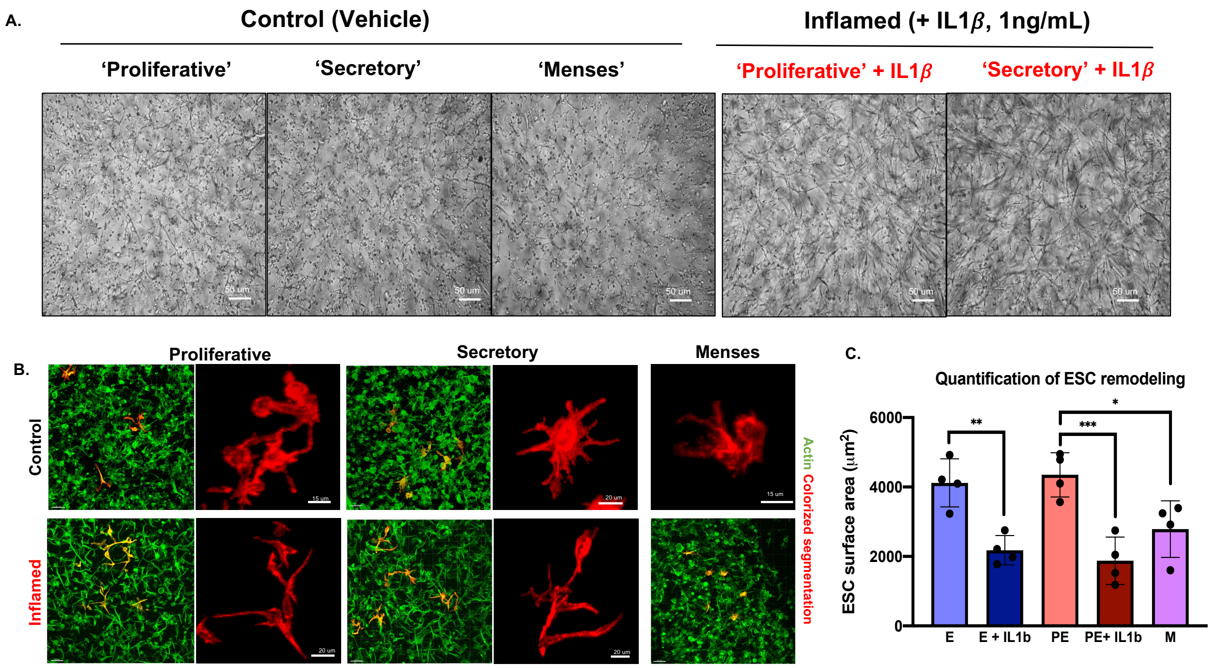

### Supplementary Fig 7. Estimating cell ratios ex vivo for tissue modeling in vitro

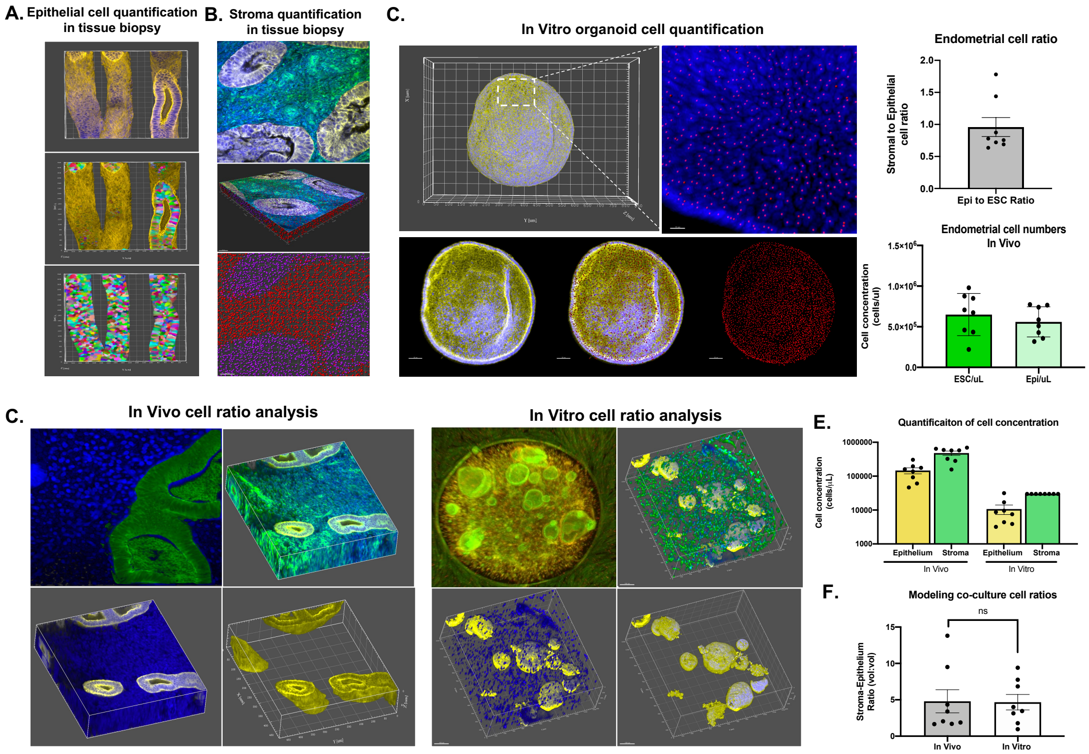

**Supplementary Fig 8. Establishment and transcriptomic analysis of monocultures and co-cocultures.**

A.

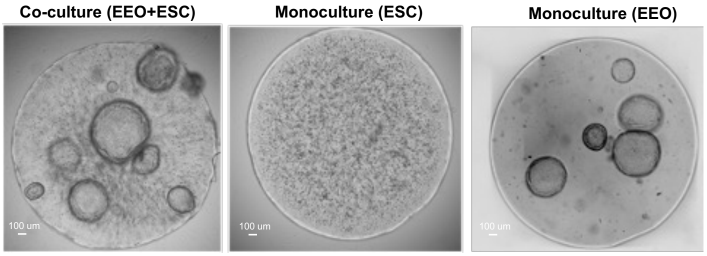


B.

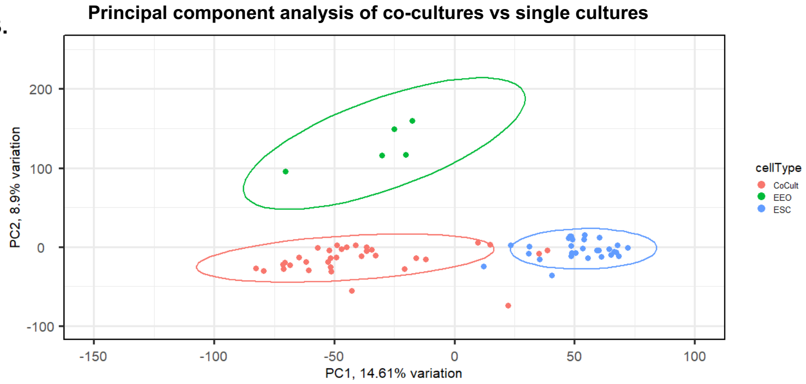


C.

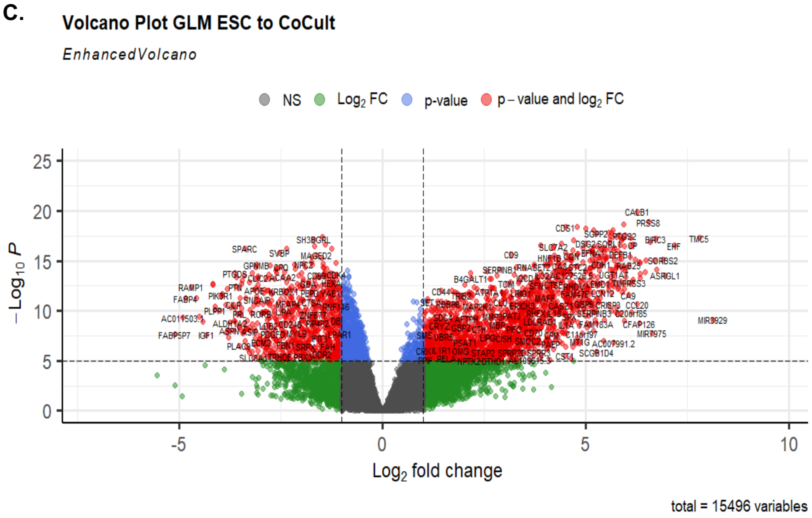

#### Supplementary Fig 9. Trancriptomic evaluation of endometrial co-culture models of the menstrual cycle phases.

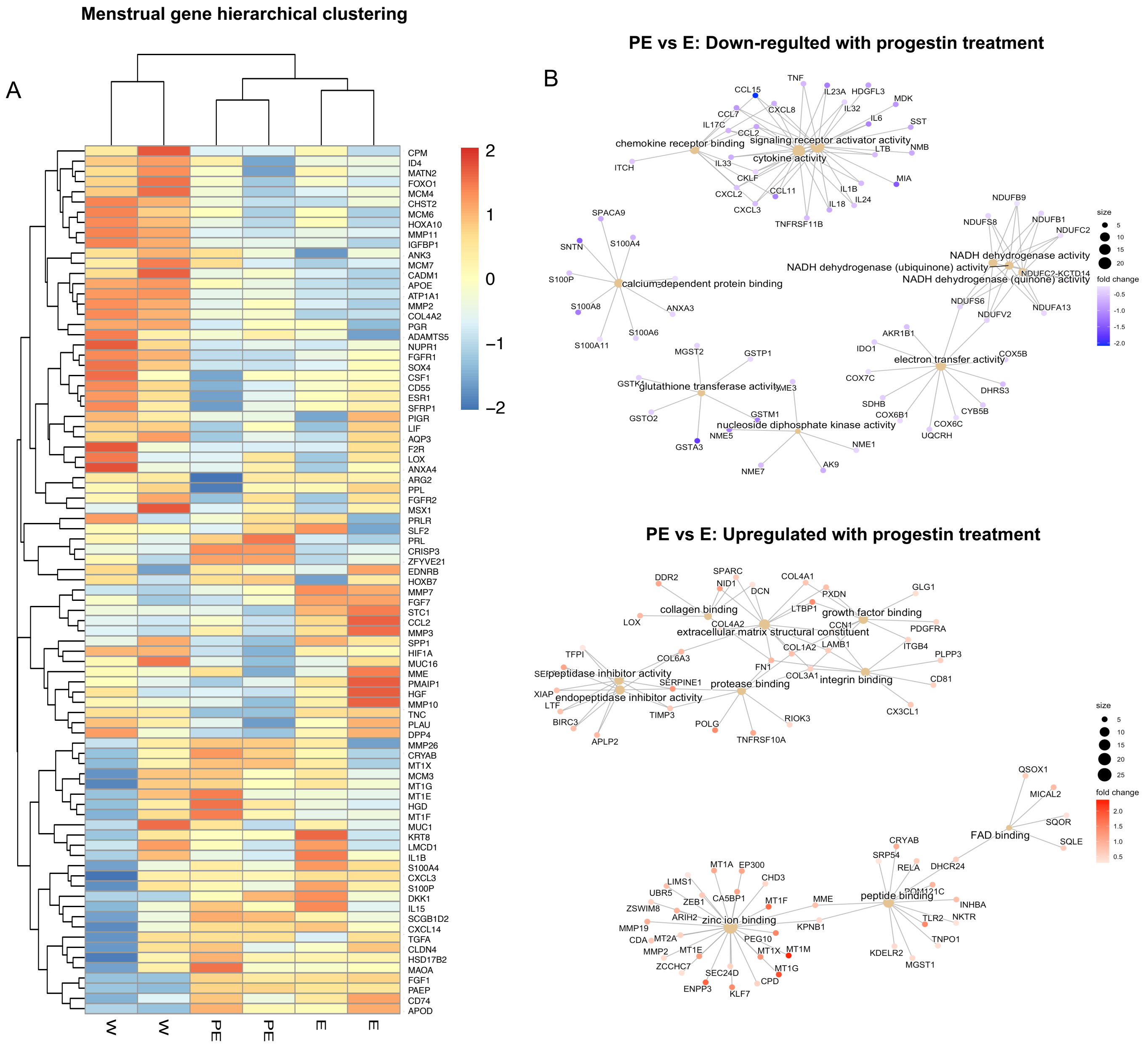

**Supplementary Fig 10. Functional and trancriptomic evaluation of endometrial co-culture recapitulation of the menstrual cycle phases.**

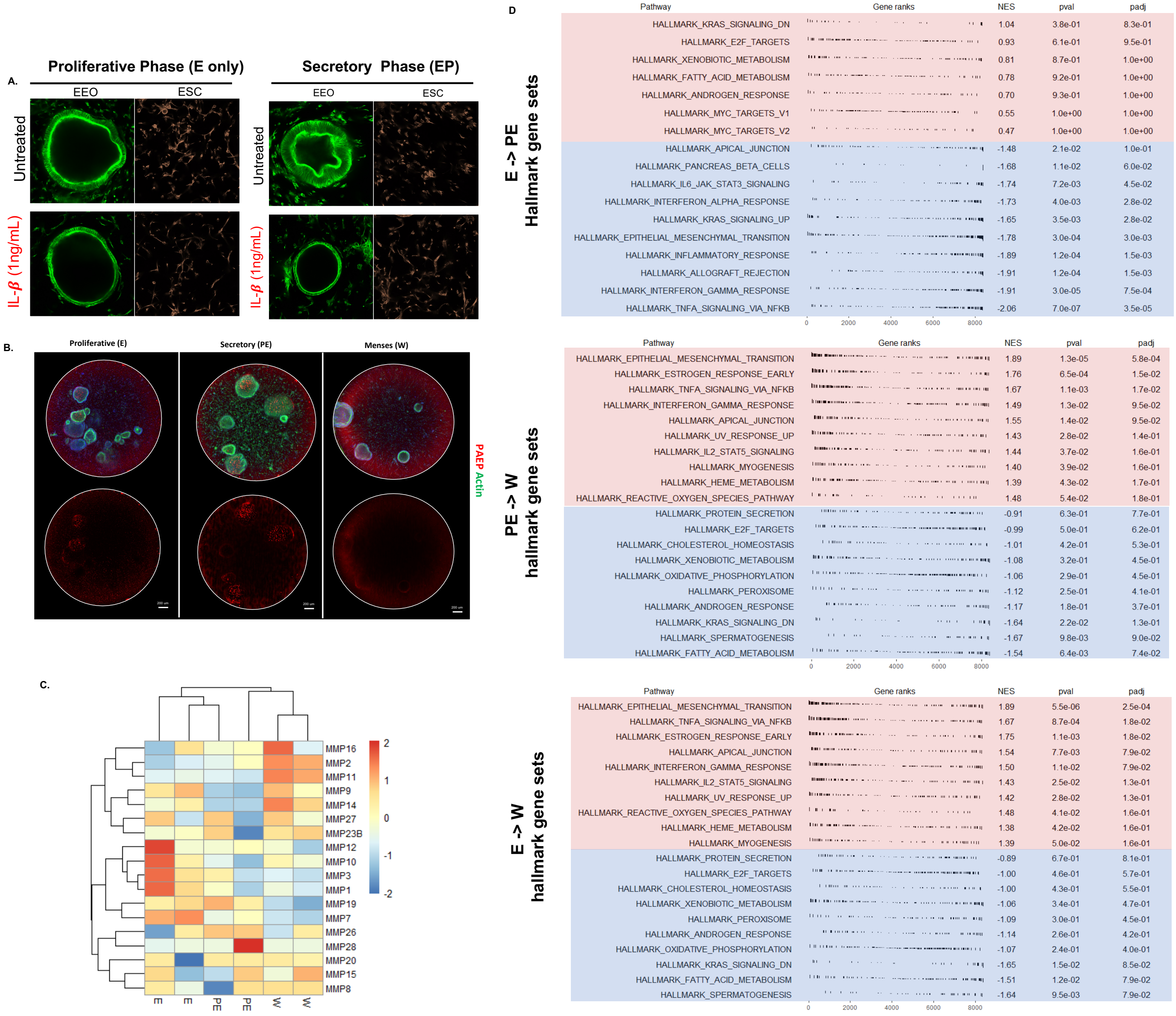

**Supplementary Fig 11. Apoptosis is increased in the menstrual phase and with IL-1β treament in the co-culture model**

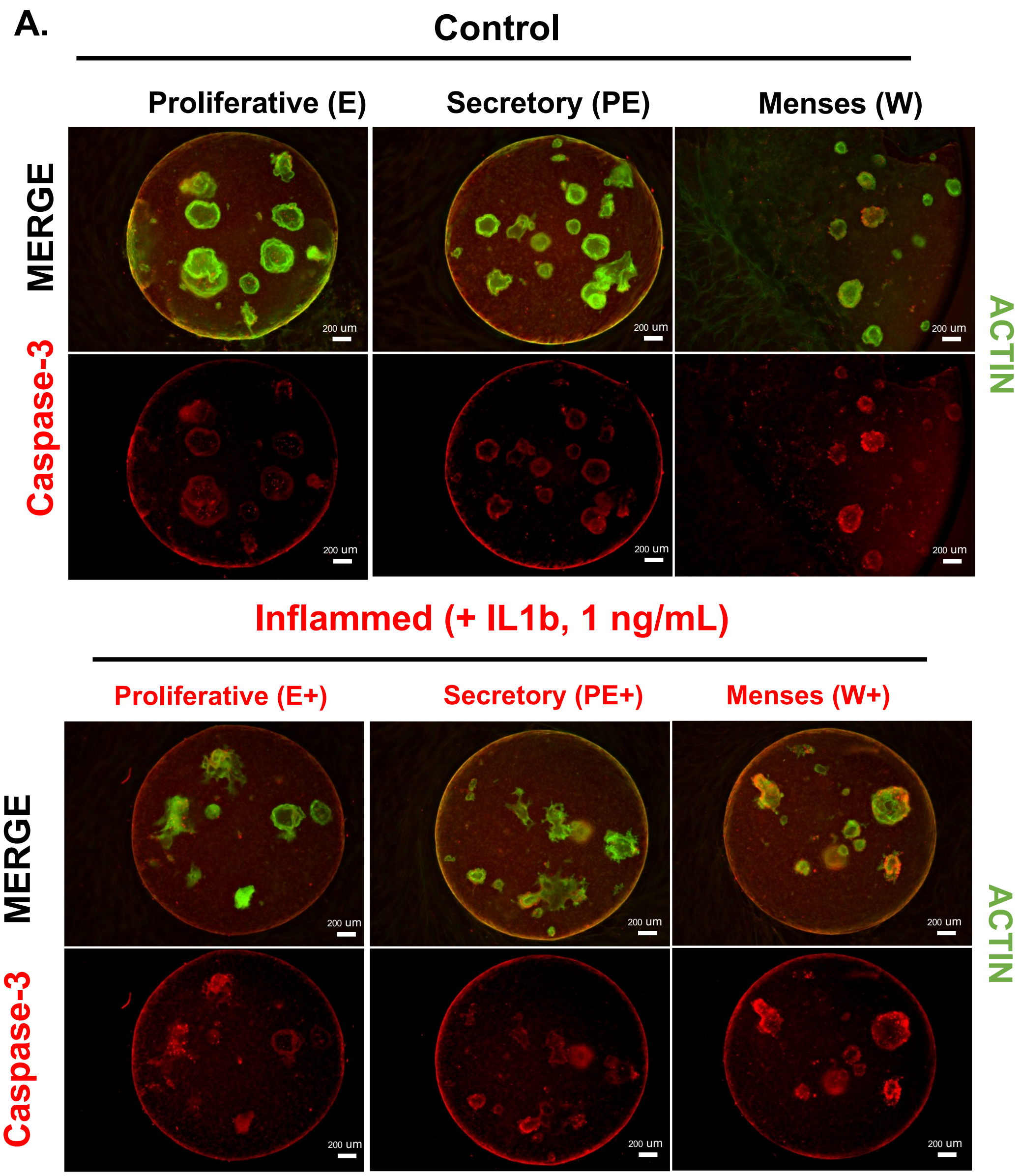

**Supplementary Fig 12. Cell viability is decreased in the IL-1β treatment group in the co-culture models at day 15 in the 3% PEG MIX hydrogel.**

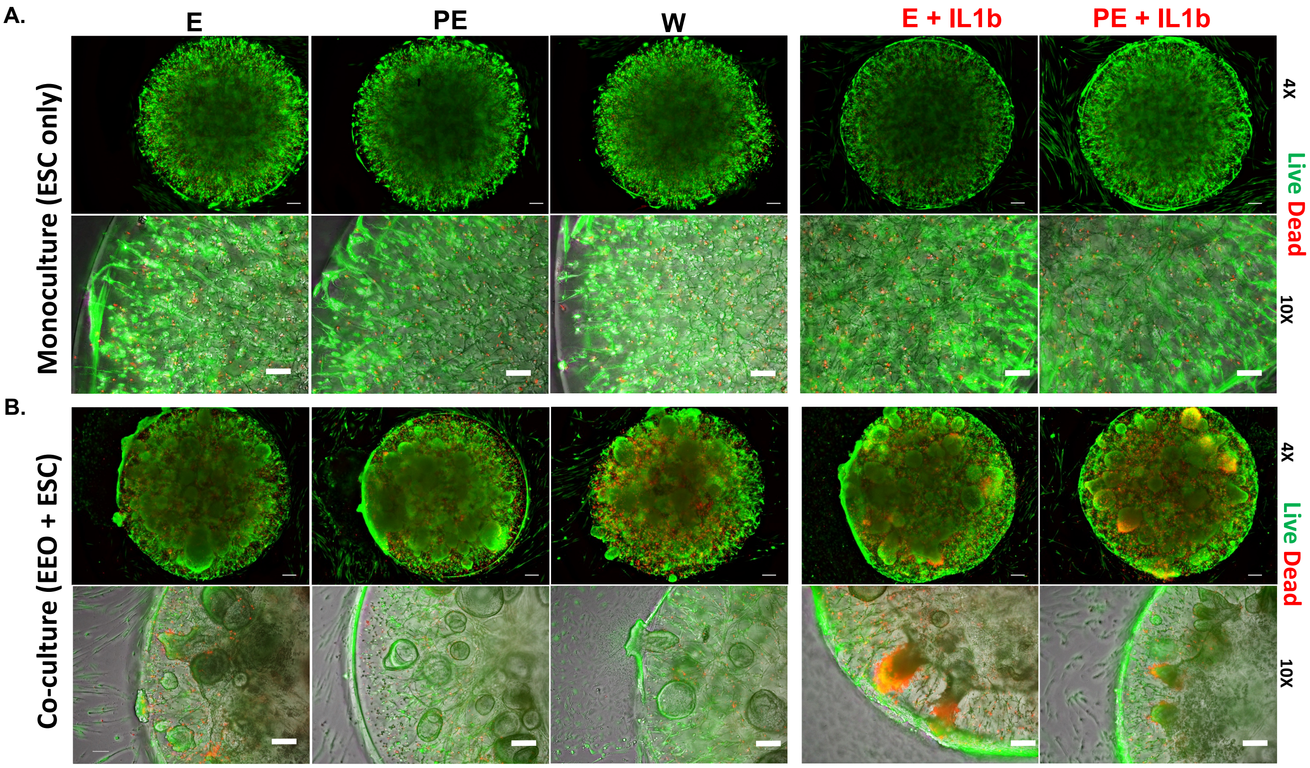

**Supplementary Fig 13. Co-cultures treated with IL-1β have transcriptomic profiles consistent with inflamed endometrial states.**

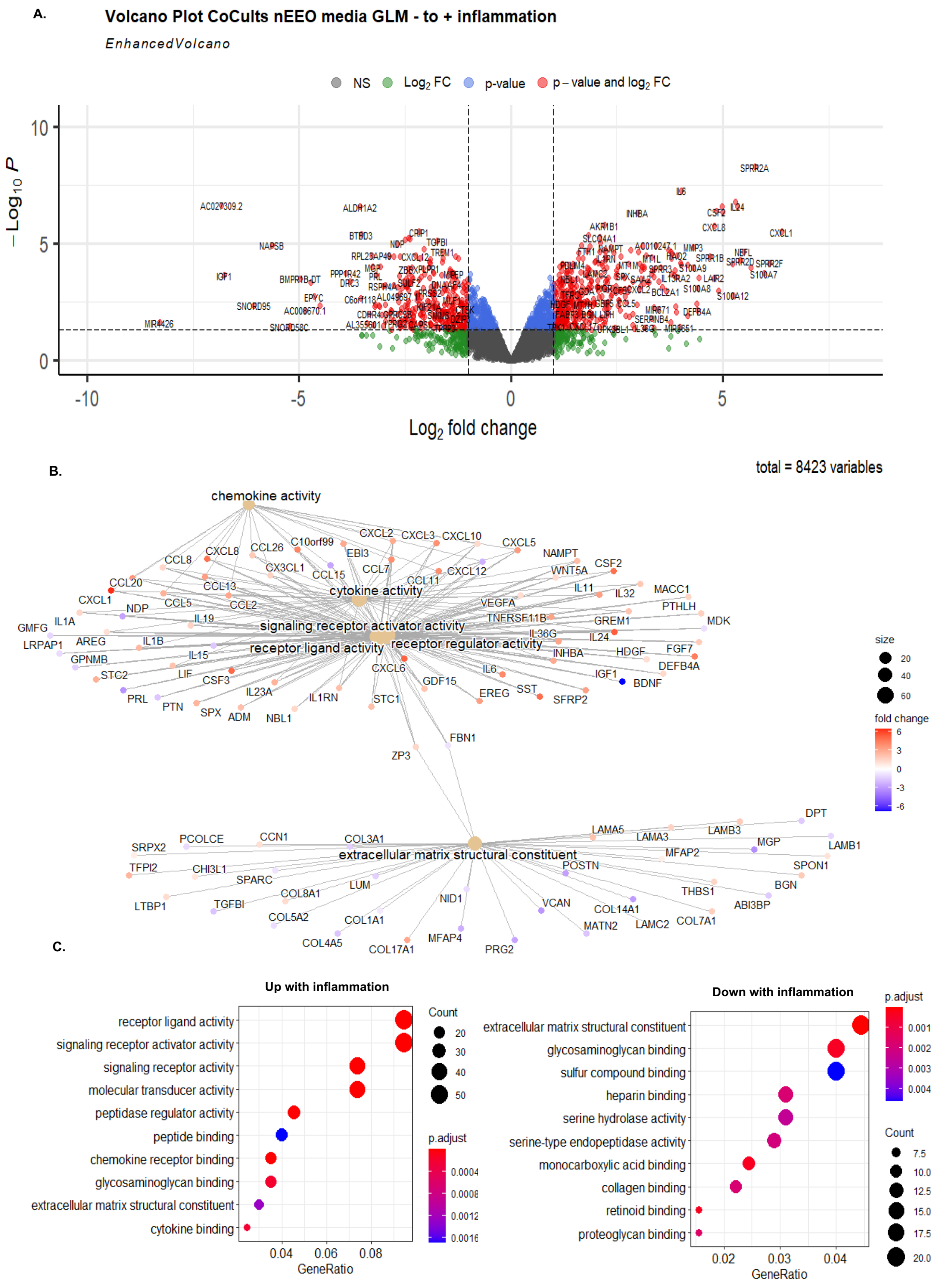

**Supplementary Fig 14. Temporal evaluation of inflammatory response of co-cultures via targeted proteomics.**

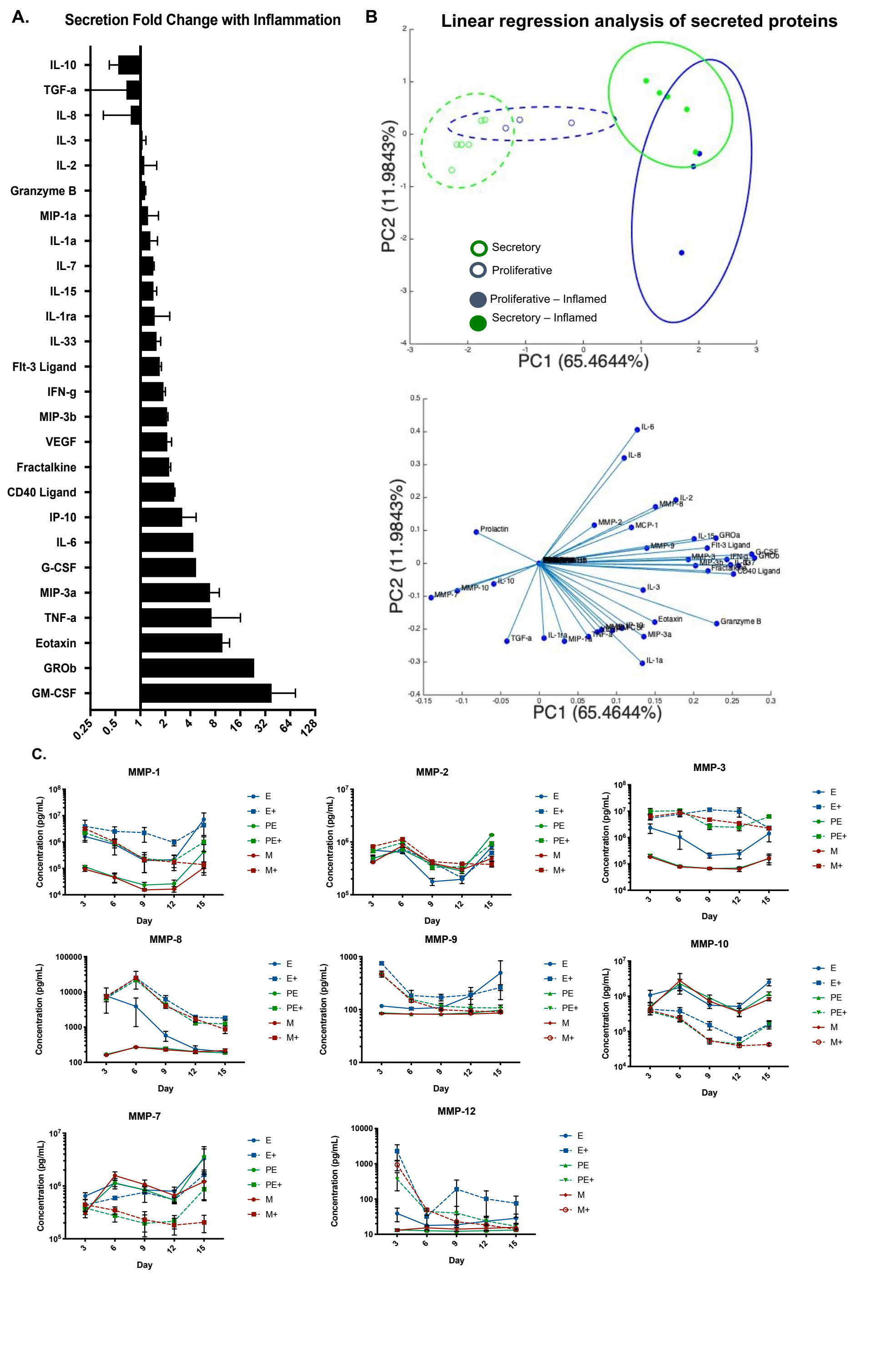

**Supplementary Fig 15. Inflammatory stimuli alter stromal morphology in endometrial co-culture.**

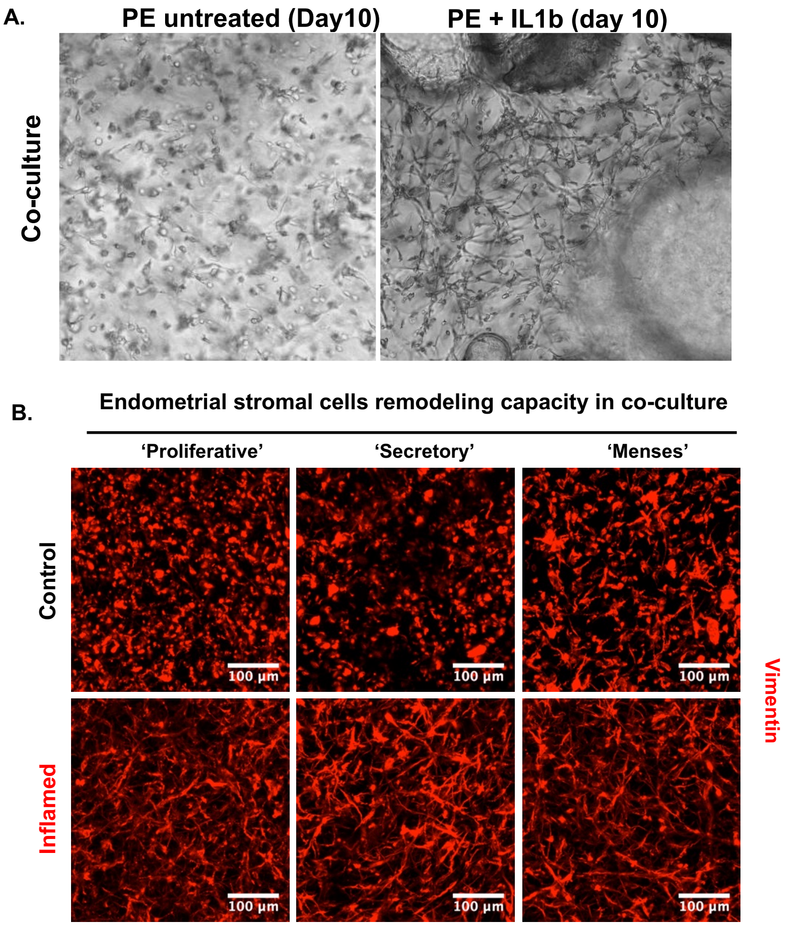

**Supplementary Fig 16. Cell number is consistent across treatment group conditions.**

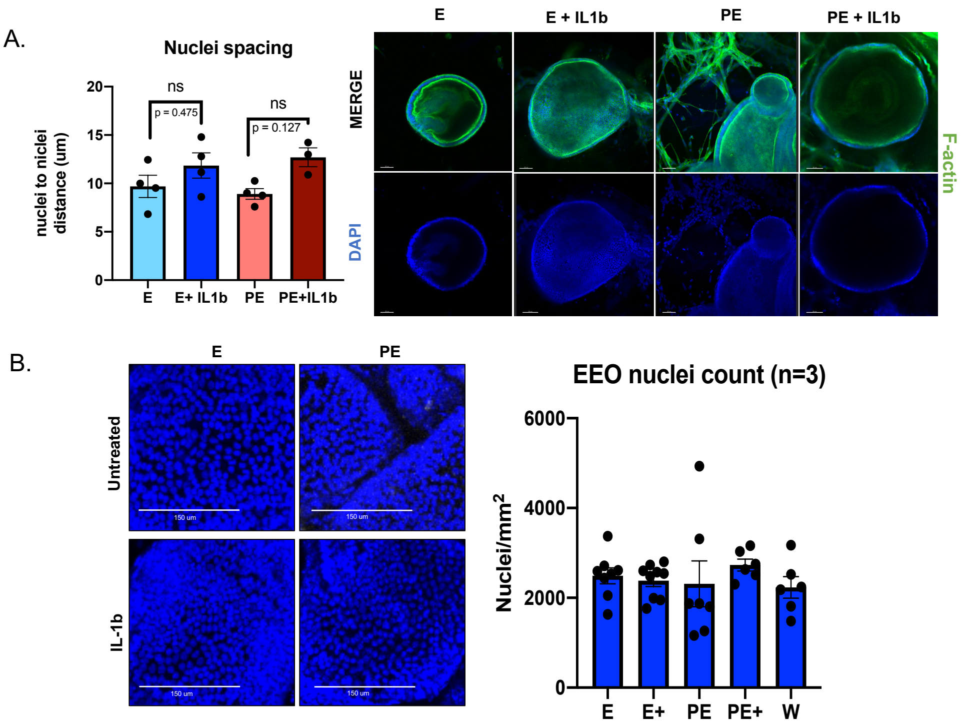

**Supplementary Fig 17. Quantitative assesment of individual EEO diamater growth rates in in co-cultures and monocultures across 15-days of culture.**

A.

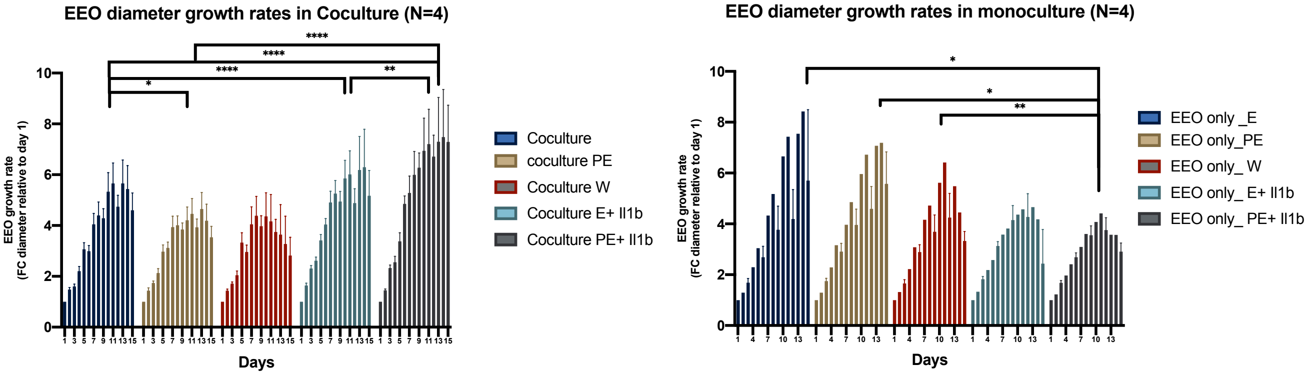


B.

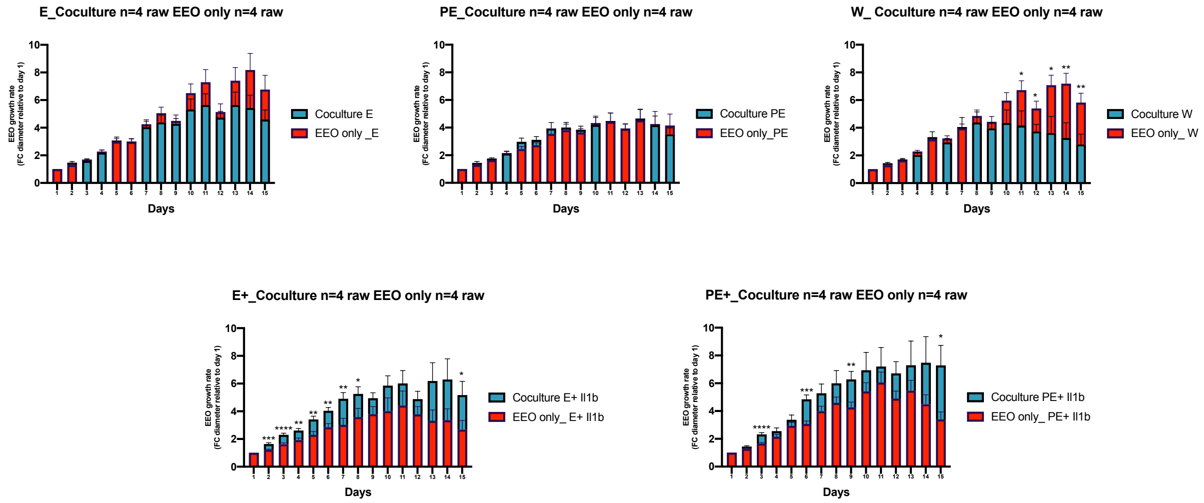

**Supplementary Fig 18. EEOs can be generated and expanded directly into synthetic ECM from primary tissues.**

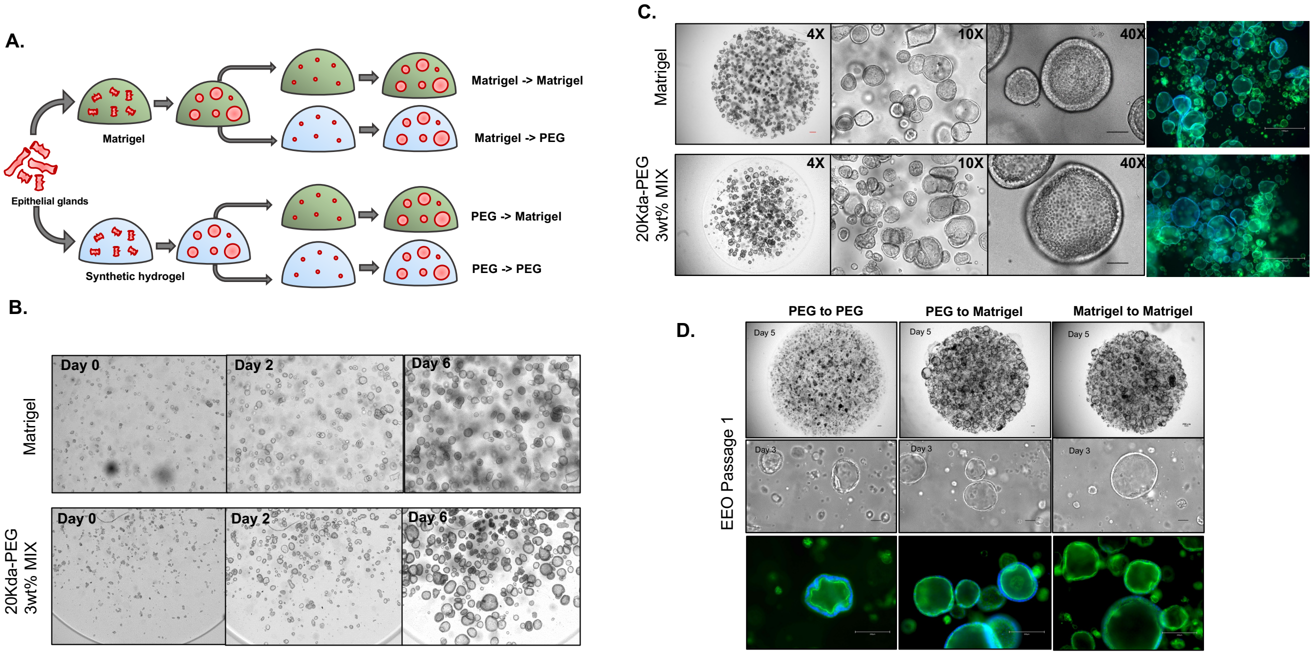

**Supplementary Fig 19. Trancriptomic evaluation of culture media effect on endometrial co-culture.**

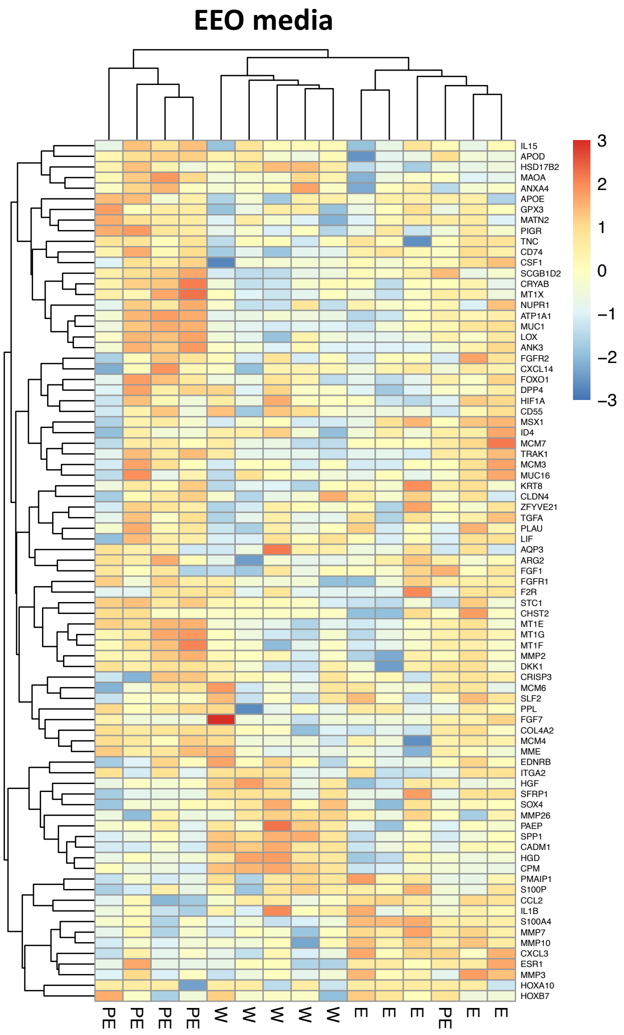

**Supplementary Fig 20. Histological characterization of endometrial tissues collected.**

A.

Donor 252 | Donor 254 | Donor 255 | Donor 259 | Donor 260

Donor 261 | Donor 262 | Donor 263 | Donor 264 | Donor 266

Donor 267 | Donor 268 | Donor 270 | Donor 271 | Donor 272

Donor 273 | Donor 274

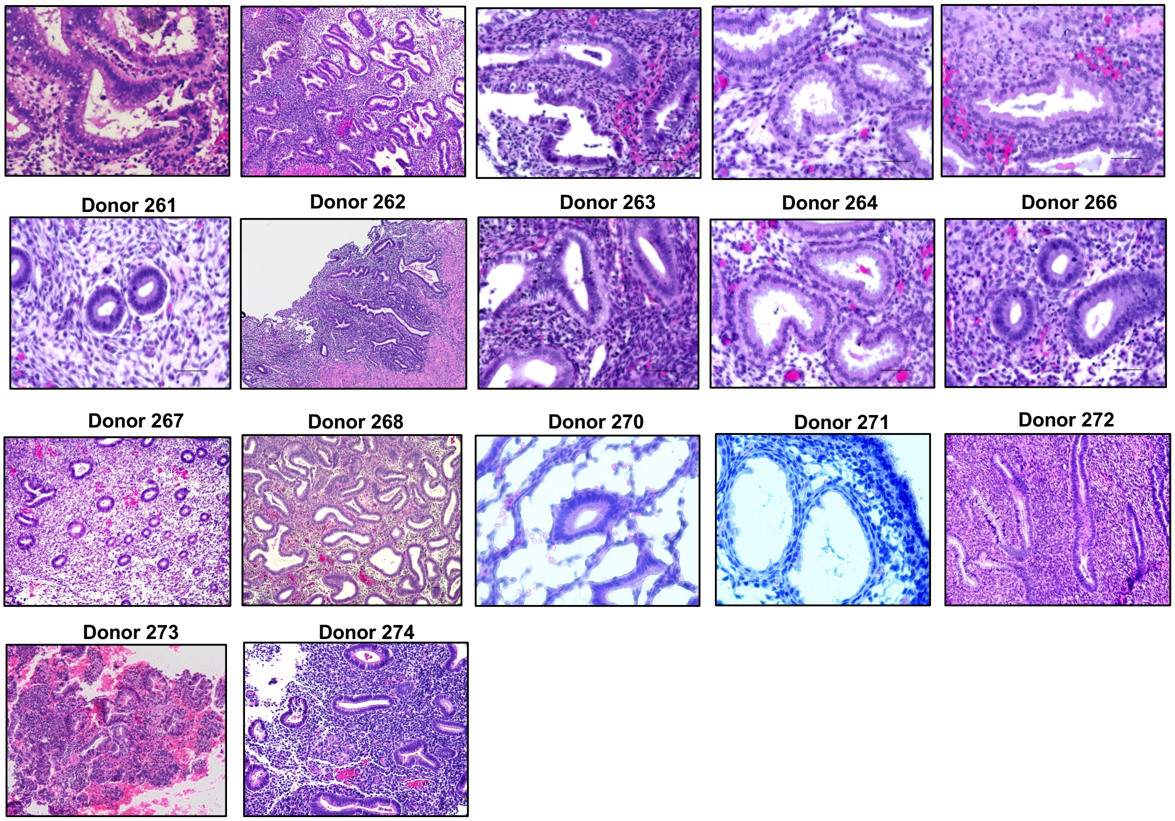

**Supplementary Fig 21. Synthetic matrix enables modeling of endometrial morphogenesis.**

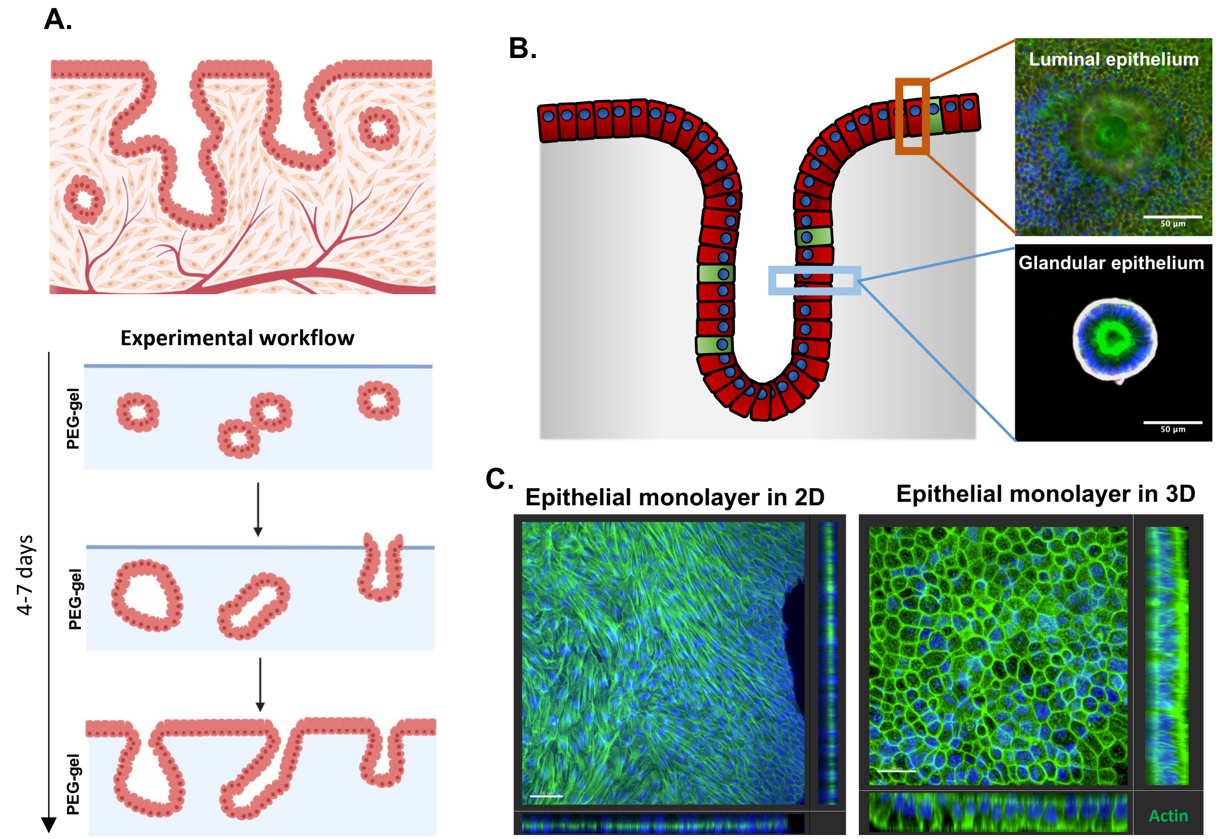
